# Galectin-3, A Novel Endogenous Trem2 Ligand, Regulates Inflammatory Response and Aβ Fibrilation in Alzheimer’s Disease

**DOI:** 10.1101/477927

**Authors:** Antonio Boza-Serrano, Rocío Ruiz, Raquel Sanchez-Varo, Yiyi Yang, Juan García-Revilla, Itzia Jimenez-Ferrer, Agnes Paulus, Malin Wennström, Anna Vilalta, David Allendorf, Jose Carlos Davila, Christopher Dunning, John Stegmayr, Sebastian Jiménez, The Netherland Brain, Maria Angustias Roca-Ceballos, Victoria Navarro-Garrido, Maria Swanberg, Christine L. Hsieh, Luis Miguel-Real, Elisabet Englund, Sara Linse, Hakon Leffler, Ulf J. Nilsson, Antonia Gutierrez, Guy C. Brown, Javier Vitorica, Jose Luis Venero, Tomas Deierborg

## Abstract

Alzheimer’s disease (AD) is a progressive neurodegenerative disease in which the formation of extracellular aggregates of amyloid beta (Aβ) peptide, intraneuronal tau neurofibrillary tangles and microglial activation are major pathological hallmarks. One of the key molecules involved in microglial activation is galectin-3 (gal3), and we demonstrate here for the first time a key role of gal3 in AD pathology. Gal3 was highly upregulated in the brains of AD patients and 5xFAD (familial Alzheimer’s disease) mice, and found specifically expressed in microglia associated with Aβ plaques. Single nucleotide polymorphisms in the *LGALS3* gene, which encodes gal3, were associated to an increased risk of AD. Gal3 deletion in 5xFAD mice attenuated microglia-associated immune responses, particularly those associated with TLR and TREM2/DAP12 signaling. *In vitro* data demonstrated the requirement of gal3 to fully activate microglia in response to fibrillar Aβ. Gal3 deletion decreased the Aβ burden in 5xFAD mice and improved cognitive behavior. Electron microscopy of gal3 in AD mice demonstrated i) a preferential expression of gal3 by plaque-associated microglia, ii) its presence in the extracellular space and iii) its association to Aβ plaques. Low concentrations (1 nM) of pure gal3 promoted cross-seeding fibrilization of pure Aβ. Importantly, a single intrahippocampal injection of gal3 along with Aβ monomers in WT mice was sufficient to induce the formation of insoluble Aβ aggregates that were absent when gal3 was lacking. High-resolution microscopy (STORM) demonstrated close co-localization of gal3 and TREM2 in microglial processes, and a direct interaction was shown by a fluorescence anisotropy assay involving the gal3 CRD domain. Furthermore, gal3 stimulated the TREM2-DAP12 signaling pathway. In conclusion, we provide evidence that gal3 is a central regulator of microglial immune response in AD. It drives proinflammatory activation and Aβ aggregation, as well as acting as an endogenous ligand to TREM2, a key receptor driving microglial response under disease conditions. Gal3 inhibition may, hence, be a potential pharmacological approach to counteract AD.

## INTRODUCTION

Alzheimer’s disease (AD) is the leading cause of dementia, affecting more than 24 million people worldwide [5] and with the incidence of AD increasing dramatically as the global population ages. The classical hallmarks of AD include the formation of amyloid-beta (Aβ) plaque deposits and of neurofibrillary tangles (NFT) containing abnormal hyperphosphorylation of tau. Over the course of the disease, the deposition of Aβ and the formation of the NFTs appear to be correlated with the on-going hippocampal neurodegeneration [7]. Hippocampal neurodegeneration has a negative effect on the cognitive ability of patients, especially short-term memory, which is impaired. The mechanisms triggering the deposition of the Aβ or the formation of NFT are currently unclear. However, several mechanisms and factors have been suggested to be involved in the initiation and the progression of the disease, including activation of the innate immune system, environmental factors and lifestyle [51]. The innate immune system has been widely studied and has been implicated in several neurodegenerative diseases [27]. Over the last years, several studies have suggested inflammation plays a major role in the initiation and progression of AD [21, 24, 28, 71, 75].

The inflammatory process in the central nervous system (CNS) is generally referred to as neuroinflammation. Glial cells have a leading role in propagating neuroinflammation. Within glial cells, microglia are considered the main source of pro-inflammatory molecules within the brain [25, 35, 72]. It is believed that sustained proinflammatory molecules such as cytokines, chemokines, nitrogen reactive species (NRS) or reactive oxygen species (ROS) can create a neurotoxic environment driving the progression of AD[1, 27, 55, 63]. Moreover, there is strong evidence supporting the role of inflammatory molecules, such as iNOS, in the initiation of plaque formation due to post-translational modifications of Aβ leading to a faster aggregation [38, 39]. Indeed, counteracting the aggregation process has been proposed as a therapeutic strategy to alleviate the progression of the pathology [67].

Conclusive evidence supporting a direct role of microglia in human neurodegeneration was revealed with the recent advent of massive genome analysis. Specifically, genetic associated studies have identified several AD risk genes strongly associated with the innate immune system, including, among others, triggering receptor on myeloid cells 2 (*trem2*), *cd33, cr1, clu, epha1, ms4a4a/ms4a6a* [8, 23]. Recent single cell transcriptomic studies of microglia have pointed out galectin-3 (Lgals3; Gal3) as one of the most attractive molecules in brain innate immunity associated with neurodegeneration (see recent review [11]). Indeed, Holtzman et al. [30], analyzed transcriptional profiles of isolated microglia from different diseases mouse models, including AD, amyotrophic lateral sclerosis and aging, and revealed a shared transcriptional network in all conditions, including a strong upregulation of gal3. Interestingly, the authors analyzed the most likely candidates to orchestrate the microglia phenotype activation and found four major hub genes: *Lgals3*, *Igf1, Csf1 and Axl* [30]. Two other recent studies characterized the molecular signature of microglial cells associated with different disease conditions including aging and AD [35, 37]. Again, a common microglia neurodegenerative disease-associated phenotype was identified, supporting that i) microglial phenotype is driven by TREM2 and ii) Gal3 was one of the most upregulated genes.

Recent findings from our group demonstrate that gal3 is released by microglia and acts as a ligand of Toll like Receptor 4 (TLR4), which is one of the canonical receptors involved in the microglia inflammatory response [10]. Based on our previous studies and recent findings from the AD field related to the role of inflammation in the disease, we hypothesize that reduction of microglial activation by removing/inhibiting gal3 will counteract the inflammatory response present in AD and slow the progression of the disease. In this work, we demonstrate i) a significant upregulation of gal3 in human AD patients compared to age-matched healthy controls, ii) the preferential expression of gal3 in microglial cells in contact with Aβ plaques both in human and mouse, iii) a clear reduction of the inflammatory response in microglial cells challenged with Aβ fibrils following gal3 inhibition and gal3 deletion in microglial cultures, iv) a significant Aβ plaque reduction and better cognitive outcome in 5xFAD mice lacking gal3, v) the release of gal3 to the extracellular space and its association to Aβ plaques, vi) the role of gal3 as a cross-seeding agent of Aβ aggregation via its carbohydrate recognition domain (CRD) at nanomolar range and vii) the modulation of microglial activation by gal3 by binding to TREM-2 via its CRD domain.

## MATERIAL AND METHODS

### Animals

5xFAD-Gal3 KO transgenic mice were generated by crossing heterozygous 5xFAD (+/-) mice with homozygous Gal3KO (-/-) mice to get 5xFAD (+/-)/Gal3 (+/-) mice. Subsequent crossings between animals expressing this genotype allowed for the generation of 5xFAD (+/-)-Gal3 (-/-) mice, hence referred to as 5xFAD/Gal3KO, and 5xFAD/Gal3 are hence referred to as 5xFAD. All animal experiments were performed in accordance with the animal research regulations (RD53/2013 and 2010/63/UE) in Spain and European Union and with the approval of the Committee of Animal Research at the University of Seville (Spain).

For primary microglial cultures, Gal3KO mice with a C57BL/6 background were obtained from Dr. K. Sävman at Gothenburg University. All procedures were carried in accordance with the international guidelines on experimental animal research and were approved by the Malmö-Lund Ethical Committee for Animal Research in Sweden (M250-11, M30-16).

### Genotyping

The genotypes of Gal3-/- (KO) and Gal3+/+ (WT) mice were determined using an integrated extraction and amplification kit (Extract-N-Amp™, Sigma-Aldrich). First, the samples were incubated at 94 °C for 5 min, followed by 40 cycles with denaturation at 94 °C for 45 sec, annealing at 55 °C for 30 sec, and elongation at 72 °C for 1.5 min. The following primers (CyberGene, Solna, Sweden) were used: galectin-3 common 5-CAC GAA CGT CTT TTG CTC TCT GG-3’), galectin-3-/- 5-GCT TTT CTG GAT TCA TCG ACT GTG G-3’ (single band of 384 bp) and galectin-3+/+ 5-TGA AAT ACT TAC CGA AAA GCT GTC TGC-3’ (single band of 300 bp) [18]. For the 5xFAD the primers (5’to 3’) used are listed below: APP Forward AGGACTGACCACTCGACCAG, APP Reverse CGGGGGTCTAGTTCTGCAT, PSN1 Forward AATAGAGAACGGCAGGAGCA, PSN1 Reverse GCCATGAGGGCACTAATCAT, WT APP Forward CTAGGCCACAGAATTGAAAGATCT, WTT APP Reverse GTAGGTGGAAATTCTAGCATCATCC, RD1, RD2 and RD3 AAGCTAGCTGCAGTAACGCCATTT ACCTGCATGTGAACCCAGTATTCTATC, CTACAGCCCCTCTCCAAGGTTTATAG. The PCR products were separated by gel electrophoresis labeled with SYBR^®^ Green (Sigma Aldrich) and visualized using a CCD camera (SONY, Tokyo, Japan).

### Protein preparation

Aβ (M1-42), i.e. with a starting Met0 to allow for production of otherwise un-tagged peptide, was expressed in *E. coli* (BL21 DE3 PLysS Star) and purified from inclusion bodies after repeated sonication using ion exchange in batch mode and size exclusion chromatography in column format, followed by lyophylization of monomer aliquots. Each day, before use in any experiment, Aβ42 monomer was again isolated from such aliquots, dissolved in 1 mL 6M GuHCl, using gel filtration (Superdex 75, 10-300 column) with 20 mM sodium phosphate, 0.2 mM EDTA, pH 8.0 as running buffer, collected on ice in a low-binding tube (Genuine Axygen Quality, Microtubes, MCT-200-L-C, Union City, CA) and the concentration was determined by absorbance at 280 nm using ε_280_ = 1440 M^-1^cm^-1^. The solution was diluted with buffer and supplemented with concentrated NaCl to achieve 10 μM monomer and 150 mM NaCl. The solution was placed in wells of a PEGylated polystyrene plate (Corning 3881) and sealed with a plastic film to avoid evaporation. Thioflavin T (ThT) was added to one well from a concentrated stock to obtain 6 μM ThT. Plates were incubated at 37 °C in a FLUOstar Omega plate reader under quiescent conditions (BMG Labtech, Offenburg, Germany) at 37 °C and the fibril formation of amyloid-beta was followed by reading the fluorescence (excitation 440 nm, emission 480 nm) through the bottom of the plate. The samples in wells without ThT were collected after reaching the plateau of the sigmoidal transition (after ca. 1 h).

### Cryogenic transmission electron microscopy (cryo-TEM)

Specimens for electron microscopy were prepared in a controlled environment vitrification system (CEVS) to ensure stable temperature and to avoid loss of solution during sample preparation. The specimens were prepared as thin liquid films, <300 nm thick, on lacey carbon filmed copper grids and plunged into liquid ethane at -180 °C. This led to vitrified specimens, avoiding component segmentation and rearrangement and water crystallization, thereby preserving original microstructures. The vitrified specimens were stored under liquid nitrogen until measured. An Oxford CT3500 cryoholder and its workstation were used to transfer the specimen into the electron microscope (Philips CM120 BioTWIN Cryo) equipped with a post-column energy filter (Gatan GIF100). The acceleration voltage was 120 kV. The images were recorded digitally with a CCD camera under low electron dose conditions. The node-to-node distance was measured using the Digital Graph software (Gatan Inc.).

### Electron microscopy immunogold labeling

12-month-old APP/PS1 mice were anesthetized with sodium pentobarbital (60 mg/kg) and transcardially perfused with 0.1 M phosphate-buffered saline (PBS), followed by 4% paraformaldehyde, 75mM lysine, 10mM sodium metaperiodate in 0.1 M phosphate buffer (PB). Vibratome 50-μm thick sections were cryoprotected in a 25% sucrose and 10% glycerol solution, followed by freezing at -80 °C in order to increase the antibody binding efficiency. Sections were then incubated 48 h in primary anti-GAL3 antibody in a PBS 0.1M/0.1% sodium azide/2% BSA-solution at 22 °C. The tissue-bound primary antibody was detected by incubation with the corresponding 1.4 nm gold-conjugated secondary antibody (1:100, Nanoprobes) overnight at 22 °C. After postfixation with 2% glutaraldehyde and washing with 50 mM sodium citrate, the labeling was enhanced with the HQ Silver™ Kit (Nanoprobes) and gold toned. Finally, the immunolabeled sections were postfixed in 1% osmium tetroxide in 0.1 M PB, block stained with uranyl acetate, dehydrated in graded acetone and flat embedded in Araldite (EMS, USA). Selected areas were cut in ultrathin sections and examined with an electron microscope (JEOL JEM 1400).

### Endotoxin test

To further evaluate the properties of our protein preparation, we performed an endotoxin assay to assure that BV2 microglial cell activation was not due to the presence of endotoxins such as LPS (Suppl. Fig 5). Fibrils preparations used for *in vitro* and *in vivo* experiments were tested for endotoxins using Pierce^®^ LAL Chromogenic Endotoxin Quantitation KIT, (ThermoScientific) according to manufacturer’s instructions (Suppl. Fig. 5A).

### XTT (cell viability) assay

XTT assays were performed to measure mitochondrial activity (mitochondrial dehydrogenase) in living cells using XTT (2,3-Bis-(2-methoxy-4-nitro-5-sulfophenyl]-2H-tetrazolium-5-carboxyanilide salt) (Sigma-Aldrich, Sweden). The assay was performed following manufacturer’s protocol in a 96-well plate (Biochrom Asys Expert 96 micro plate reader, Cambridge, UK) (Suppl. Fig. 5B).

### In vitro experiments. Cell lines and primary cultures

BV2 microglial cells were cultured in 12-well plates, 250.000 cells/well and stimulated with Aβ monomers and fibrils for a range of time points (3, 6, 12 and 24 h) and concentrations (3 and 10 μM). Primary microglial cells were obtained from WT and Gal3KO mice. The primary cells were obtained from the cortex, as previously described [16], and cultured for 14 days in T75 flask culture conditions before treatment. After 14 days, microglial cells were isolated and 25,000 were cultured in 96-well plates and incubated for 12 h with Aβ fibrils. Cell viability and number of cells were assessed using a TC20 Bio-Rad Cell counter, Bio-Rad chambers slides and Trypan Blue 0.04% for BV2 and primary cultures. The same number of viable cells of each genotype (WT or Gal3KO) was plated per well. The cells were grown in DMEM (Invitrogen), FBS (Invitrogen) 10 % (v/v) and Penicillin-Streptomycin (Invitrogen) 1 % (v/v) in 5 % CO2 in air at 37 °C in a humidified incubator. After incubation, the medium was collected to measure extracellular cytokines and intracellular proteins in order to evaluate the microglial activation profile.

### DAP12-TREM-2 reporter cell line

The ability of gal3 to activate TREM2-DAP12 signaling was assayed in a BWZ thymoma reporter cell line transfected with TREM2 and DAP12 as previously described [31]. In these cells, TREM-2/DAP12 signaling activates phospholipase C, leading to a calcium influx that activates calcineurin, leading to disinhibition of the nuclear import of NFAT (nuclear factor of activated T-cells), which induces transcription of the LacZ β-galactosidase gene. The TREM2/DAP12 reporter cells or parental BWZ cells not expressing TREM2 or DAP12 (control) were incubated with the indicated concentrations of gal3 for 24 h at 37 °C, washed, and lysed in a buffer containing 100 mM 2-ME, 9 mM MgCl2, 0.125% NP-40, and 30 mM chlorophenol red galactosidase (CPRG). Plates were developed for 24 h at 37 °C, and lacZ activity was measured as previously described [31]. As a positive control, ionomycin (3 μM) was added.

### Phagocytosis experiments

After 24h of primary microglia culturing, gal3 was added at 1μM concentration for 30 minutes prior to incubation with labeled Aβ1-42 (HiLyte™ Fluor 647-labeled, Human; ANA64161) for one hour. Aβ1-42 was added as fibrillar. Fibrillar Aβ1-42 samples were obtained by incubating them for 24 hours at 37 °C. Cells were detached by brief incubation with trypsin with EDTA and analyzed by flow cytometry (Accuri C6, BD Biosciences). Mean fluorescence in FL4 gate was used to plot results and expressed as percentage from Aβ1-42 uptake into primary microglia.

### Sequential protein extraction

Soluble and insoluble protein fractions were obtained from the whole cortex (mice) and temporal cortex (human) using sequential protein extraction. Fractions were obtained by disruption of the cortex with a dounce homogenizer in the presence of PBS (1 mL/100 μg of tissue). Supernatants, S1 fractions, were aliquoted and stored at -80 °C. S1 soluble fraction was obtained after centrifugation for 1 h at 40,000 rpm in special tubes for high-speed centrifugation by Beckman-Coulter. The pellets were extracted in RIPA buffer (Sigma-Aldrich, Germany) ultracentrifuged at 30,000 rpm and supernatants, S2 fractions (intracellular particulate proteins), were aliquoted and stored. Pellets were re-extracted in buffered-SDS (2% SDS in 20 mM Tris–HCl, pH 7.4, 140 mM NaCl), centrifuged as above and supernatants, S3 (SDS releasable proteins) were stored. Finally, the remaining pellets (P3) were extracted in SDS-urea (20 mM Tris–HCl, pH 7.4, 4% SDS and 8 M urea). PBS and RIPA solution were prepared using a protein inhibitor (Protein Inhibitor Cocktail, ThermoScientific) to prevent protein degradation and to inhibit the enzymatic activity of phosphatases (PhosphoStop, Roche).

### Western blot

Proteins were extracted from cell cultures with RIPA buffer (Sigma-Aldrich, Germany) along with proteinase and phosphatase inhibitors (Roche, Switzerland). Protein concentration was measured using a BCA kit according to manufacturer’s protocol (BCA Protein Assay-Kit, ThermoScientific, Sweden). Protein extracts were then separated by SDS-PAGE using pre-cast gels (4-20% from Bio-Rad) in TGS buffer (Bio-Rad, Sweden). The proteins were transferred to nitrocellulose membranes (Bio-Rad, Sweden) using the TransBlot turbo system from Bio-Rad. The membranes were subsequently blocked for 1 h with skim milk at 3% (w/v) in PBS, then washed 3×10 min in PBS supplemented with 0.1% (w/v) Tween 20 (PBS-T). Then, blots were incubated with primary antibodies in PBS-T, as mentioned above, overnight. Following this, we incubated with secondary antibodies for 2 h. After the secondary antibody, we washed 3 times with PBS-T, and then blots were developed using ECL Clarity (Bio-Rad) according to the manufacturer’s protocol and using the ChemiBlot XRS+ system from Bio-Rad.

### ELISA Plates

MSD (MesoScale) plates were used to evaluate the cytokine (proinflammatory panels for IFN-γ, IL-1β, IL-2, IL-4, IL-5, IL-6, IL-8, IL-10, IL-12 and TNF-α) levels in culture medium, blood and brain soluble fraction from both human and mouse samples. To measure the cytokines in mouse and human soluble fractions, we pulled together 50 μg of the S1 and S2 fraction. MSD plates were also used to measure the levels of Aβ38/40/42, phosphorylated Tau and Total Tau in the soluble and insoluble fraction of the brain extraction from WT, Gal3 KO, 5xFAD and 5xFAD Gal3 KO mice. Serial dilutions of the soluble and the insoluble fraction were tested in order to obtain an accurate measure of the protein levels. 1 μg from the soluble fraction was diluted to evaluate Aβ38/40/42, and 0.3 μg from the insoluble fraction was diluted to evaluate Aβ38/40/42. The plates were developed using the 4x reading buffer dilute to a factor of 1x with distilled water, and the plates were read using the QuickPlex Q120 reader from Mesoscale. The detection ranges of the different cytokines measured were as follows: IL1β (1670-0,408 pg/mL), IL4 (1660-0,405 pg/mL), IL12 (32200-7,86), IL10 (3410-0,833), IFN-γ (938-0,229 pg/mL), IL2 (2630-0,642 pg/mL), IL5 (967-0,236 pg/mL), IL6 (5720-1,40), KC/GRO (1980-0,483 pg/mL) and TNF-α (627-0,153). Aβ_40_ detection range was (15100-3,69 pg/mL), and Aβ_42_ detection range was (2280-0.557 pg/mL).

### Immunohistochemistry and immunofluorescence

#### Immunohistochemistry

Mice were transcardially perfused under deep anesthesia with 4% paraformaldehyde and PBS, pH 7.4. The brains were removed, cryoprotected in sucrose and frozen in isopentane at −15 °C, after which were cut in 40 μm thick slices in the coronal plane on a freezing microtome and serially collected in wells containing cold PBS and 0.02% sodium azide. 5xFAD and 5xFAD/Gal3KO free-floating sections were first treated with 3% H_2_O_2_/ 10% methanol in PBS, pH 7.4 for 20 min to inhibit endogenous peroxidases and then with avidin-biotin Blocking Kit (Vector Labs, Burlingame, CA, USA) for 30 min to block endogenous avidin, biotin and biotin-binding proteins. For single immunolabeling, sections were immunoreacted with one of the primary antibodies: anti-Aβ42 rabbit polyclonal (1:5000 dilution, Abcam), anti-amyloid precursor protein (APP) rabbit polyclonal (1:20000 dilution, Sigma), anti-CD45 rat monoclonal (clone IBL-3/16, 1:500 dilution, AbD Serotec), anti-galectin3 (gal3) goat polyclonal (1:3000 dilution, R&D) over 24 or 48 h at room temperature. The tissue-bound primary antibody was detected by incubating for 1 h with the corresponding biotinylated secondary antibody (1:500 dilution, Vector Laboratories), and then followed by incubating for 90 min with streptavidin-conjugated horseradish peroxidase (Sigma–Aldrich) diluted 1:2000. The peroxidase reaction was visualized with 0.05% 3-3-diaminobenzidine tetrahydrochloride (DAB, Sigma Aldrich), 0.03% nickel ammonium sulphate and 0.01% hydrogen peroxide in PBS. After DAB, some sections immunolabeled for APP, CD45 or Gal3 were incubated 3 min in a solution of 0.2% of Congo red. Sections were mounted on gelatine-coated slides, air-dried, dehydrated in graded ethanol, cleared in xylene cover with DPX (BDH) mounting medium. Omitting the primary antisera controlled for specificity of the immune reactions.

#### Immunofluorescence

For double CD45/Gal3 or Iba1/Gal3 immunofluorescence free-floating sections were first incubated with the primary antibodies followed by the corresponding Alexa 488/568 secondary antibodies (1:1000 dilution, Invitrogen). Sections were embedded in autofluorescence eliminator reagent (Merck Millipore), following the manufacturer’s recommendations, to eliminate fluorescence emitted by intracellular lipofuscin accumulation. Finally, sections were coverslipped with 0.01M PBS containing 50% glycerin and 3% triethylenediamine and examined under a confocal laser microscope (Leica SP5 II) or Olympus BX-61 epifluorescent microscope.

For Aβ (1:1000, Sigma-Aldrich), gal3 (1:1000, R&D Systems), TREM2 (1:500, R&D) and Iba 1 (1:500, WAKO) tissue was incubated for 24 h and the following day, the brain sections were rinsed for 1 h in PBS containing 0.1% Triton X-100. After incubating for 1h with the corresponding secondary antibodies (1:500, Alexa antibodies, Invitrogen), and rinsing again with PBS containing 0.1% Triton X-100 for 60 min. Then, brain sections were mounted in glycerol 50% for visualization. Fixed tissue was examined in an inverted ZEISS LSM 7 DUO confocal laser-scanning microscope using a 20x air objective with a numerical aperture of 0.5. All Images were obtained under similar conditions (laser intensities and photomultiplier voltages), and usually on the same day. Morphometric analysis of the fluorescently labeled structures was performed offline with Fiji ImageJ software (W. Rasband, National Institutes of Health). Areas for the specific antibodies were determined automatically by defining outline masks based on brightness thresholds from maximal projected confocal images [4].

### Plaque loading quantification

Plaque loading was defined as the percentage of hippocampal CA1 region stained for Aβ (with anti Aβ42) Quantification of extracellular Aβ content neurites content was done as previously described [60, 64]. Images were acquired with a Nikon DS-5M high-resolution digital camera connected to a Nikon Eclipse 80i microscope. The camera settings were adjusted at the start of the experiment and maintained for uniformity. Digital 4x (plaques) images from 6- and 18-month-old 5xFAD and 5xFAD-Gal3KO mice (4 sections/mouse; n = 6-7/age/genotype) were analyzed using Visilog 6.3 analysis program (Noesis, France). The hippocampal area (CA1 or Thalamus) in each image was manually outlined, leaving out pyramidal and granular layers in the case of APP quantification. Then, plaque areas within the hippocampal regions were identified by level threshold that was maintained throughout the experiment for uniformity. The color images were converted to binary images with plaques. The loading (%) for each transgenic mouse was estimated and defined as (sum plaque area measured/sum hippocampal area analyzed) × 100. The sums were taken over all slides sampled and a single burden was computed for each mouse. The mean and standard deviation (S.D.) of the loadings were determined using all the available data. Quantitative comparisons were carried out on sections processed at the same time with same batches of solutions.

### In vitro assay for measuring Aβ 1-42 aggregation

Human amyloid-beta 1-42 (Anaspec) was prepared as previously described [52]. Briefly, the peptide was dissolved in 1,1,1,3,3,3-hexafluoroisopropanol (HFIP, Sigma) and dried under a stream of nitrogen. Prior to the aggregation assay, Aβ 1-42 was resolubilized by DMSO (Sigma) and resuspended in aggregation medium DMEM/F12. Aβ aggregation was measured by thioflavin T fluorescence as previously described [50]. Briefly, thioflavin T (Sigma) and Aβ (10 μM final concentration for both) were added to the physiological growth medium DMEM/F12 in the presence or absence of gal3 at 1 nM. Aβ was allowed to aggregate for ~15 h at 31.5 °C and fluorescence (440 nm absorbance, 480 nm emission) was measured in 10 min intervals by a spectophotometer (FLUOstar OPTIMA, BMG Labtech). ThT fluorescence was normalized to initial fluorescence values (10-30 min after start of aggregation). Aggregates in wells were imaged using a fluorescence microscope (DMI6000; Leica) using 480/40 nm excitation and 527/30 emission filters with a 10x objective. To assess the source of fluorescence amyloid aggregates in medium were centrifuged for 15 min at 15,000g and their fluorescence was re-measured.

### Hippocampal Aβ injections in wild-type mice

Aβ monomers (10 μM) and Aβ monomers together with gal3 (10 μM) were pre-incubated for 1 h at 37 °C prior to the intracerebral injections. A volume of 2 ul was injected (0.5 ul per min) in the dentate gyrus. In the left hemisphere, we injected only Aβ monomers and in the right hemisphere Aβ monomers with gal3. Two months after injections, the mice were sacrificed and transcardially perfused in 4% PFA and postfixed in 4% PFA overnight followed by sucrose saturation (25% in PBS). Brain were sectioned by a microtome (30 μm) and stained for ThioS, 6E10, GFAP, NeuN and Iba1.

### Stochastic optical reconstruction microscopy (STORM)

The samples were the same that we used for the confocal images (see the protocol in immunofluorescence staining section in material and methods) except we changed the buffer and the detection device. Images were acquired as previously described by Van der Zwaag et al [65]. Briefly, images were acquired using a Nikon N-STORM system configured for total internal reflection fluorescence (TIRF) imaging. STORM buffer contains 10 mM Tris pH 8, 50 mM NaCl, oxygen scavenging system (0.5 mg/mL glucose oxidase (Sigma-Aldrich), 34 μg/mL catalase (Sigma), 5% (w/v) glucose and 100 mM cysteamine (Sigma-Aldrich). Excitation inclination was tuned to adjust the focus and to maximize the signal-to-noise ratio. Fluorophores were excited illuminating the sample with the 647 nm (~ 125 mW), and 488 nm (~50 mW) laser lines built into the microscope. Fluorescence was collected by means of a Nikon APO TIRF 100x/1.49 Oil W.D. 0.12 mm. Images were recorded onto a 256 × 256 pixel region of a EMCCD camera (Andor Ixon3 897). Single molecule localization movies were analyzed with NIS element Nikon software.

### Immunofluorescence of human sections

Endogenous peroxidases were deactivated by incubation in peroxidase block for 15 min with gentle agitation. The sections were then washed (3 × 15 min) in 0.1 M KPBS, after which they were incubated in blocking buffer (5% goat serum blocking with 0.1 M KPBS and 0.025% Triton-X) for at least 1 h with gentle agitation. The sections were then washed (3 × 15 mins) in 0.1 M KPBS, and the primary antibody added (1:300). The sections were then incubated at 4 °C overnight with gentle agitation. The sections were washed (3 × 15 min) in 0.1 M KPBS, after which poly-HRP secondary antibody was added. The sections were then incubated for 1 h at room temperature. For triple Iba1/Gal3/Amyloid-beta immunofluorescence, sections were first incubated with the primary antibodies followed by the corresponding Alexa 647/488/555 secondary antibodies (1:1000 dilution, AlexaFluor, Life Technologies). Sections were embedded in 0.6 g Sudden Black (Sigma) dissolved in 70% ethanol. The sections, after mounting and drying on slide, were incubated in the sudden black solution for 5 min. Thereafter, the sections were washed in PBS and mounted with mounting medium. The camera settings were adjusted at the start of the experiment and maintained for uniformity. Confocal Microscope Nikon Eclipse Ti (Nikon, Japan) and NIS elements software (Nikon, Japan) were used to take 20x magnification pictures and for the final collage.

### Fluorescent anisotropy

#### Production of recombinant human galectins

Recombinant human galectins (*i.e.* Gal3 wild type and Gal3 R186S mutant) were produced in *E. Coli* BL21 Star (DE3) cells and purified by affinity chromatography on lactosyl-sepharose columns, which has been previously described by Salomonsson et al. [58].

#### Establishment of the affinity between galectins and TREM2

A fluorescence anisotropy (FA) assay was used to determine the affinity of recombinant TREM2 and wild type or mutant Gal3 in solution. The method has previously been described in detail by Sörme et al. for saccharides and synthetic small-molecule galectin inhibitors [62]. In short, increasing concentrations of galectins were first titered against a fixed concentration of saccharide probe (0.02 μM), and, when this is done, the anisotropy value increases from a value for probe free in solution (A_0_) to a value where all probe molecules are bound to galectins (A_max_). To establish the dissociation constant (K_d_) values between TREM2 and the Gal3 wild type or R186S mutant, a competitive variant of the FA assay was used. In this assay, increasing concentrations of TREM2 were titered against fixed concentrations of galectin and probe (see below for details). By obtaining the anisotropy values for the different TREM2 concentrations, together with the values for A_max_ and A_0_, the K_d_ values could be calculated according to the equations presented in Sörme et al. [62].

The FA of the fluorescein-conjugated probes was measured using a PheraStarFS plate reader and PHERAstar Mars version 2.10 R3 software (BMG, Offenburg, Germany). The excitation wavelength used was 485 nm, and the emission was read at 520 nm. All experiments were performed in PBS at room temperature (~20 °C). The anisotropy values for each data point were read in duplicate wells of 386-well plates (at a total volume of 20 μl). K_d_ values were calculated as weighted mean values from concentrations of TREM2 that generated between 20-80% inhibition (where inhibition values of approximately 50% had the highest impact on the mean value).

*Gal3 (wild type) affinities:* experiments were performed with Gal3 at a concentration of 0.30 μM and the fluorescent probe 3,3’-dideoxy-3-[4-(fluorescein-5-yl-carbonylaminomethyl)-1H-1,2,3-triazol-1-yl]-3’-(3,5-dimethoxybenzamido)-1,1’-sulfanediyl-di-β-d-galactopyranoside at 0.02 μM[59].

*Gal3 (R186S mutant) affinities:* experiments were performed with Gal3 R186S at a concentration of 2 μM and the fluorescent probe 2-(fluorescein-5/6-yl-carbonyl)aminoethyl-2-acetamido-2-deoxy-α-d-galactopyranosyl-(1–3)-[α-l-fucopyranosyl-(1–2)]-β-d-galactopyranosyl-(1–4)-β-d-glucopyranoside at 0.02 μM[14].

### TREM2 and galectin-3 3D modeling

Extracellular domain of TREM2 (white) with tetratennary N-glycan (stick-model) and Gal3 CRD (yellow) modeled in. Mutations in TREM2 with increased risk for AD are blue and for Nasu-Hakola disease (NHD) are red. The C-terminus of the extracellular fragment that normally links further to the transmembrane domain is green.

The TREM2 model is from pdb 5 ELI, published in Kober DL, Alexander-Brett JM, Karch CM, Cruchaga C, Colonna M, Holtzman MJ, Brett TJ. Neurodegenerative disease mutations in TREM2 reveal a functional surface and distinct loss-of-function mechanisms. Elife. 2016 Dec 20;5. pii: e20391. doi: 10.7554/eLife.20391.

A tetranatennary N-glycan was modeled in at the single N-glycosylation site using the GlyCam server, Woods Group. (2005-2018) GLYCAM Web. Complex Carbohydrate Research Center, University of Georgia, Athens, GA. (http://glycam.org). The model of bound Gal3 CRD was from a structure in complex with LacNAc (pdb IKJL, Sorme P, Arnoux P, Kahl-Knutsson B, Leffler H, Rini JM, Nilsson UJ Structural and thermodynamic studies on cation-Pi interactions in lectin-ligand complexes: high-affinity Gal3 inhibitors through fine-tuning of an arginine-arene interaction J. Am. Chem. Soc. (2005) 127 p.1737-1743), which was superimposed on the terminal LacNAc of the N-glycan. The picture was made with The PyMOL Molecular Graphics System, Version 2.0 Schrödinger, LLC.

### Gene Array

Hippocampal samples from 5xFAD, 5xFAD/Gal3KO, WT and Gal3KO mice at 6 and 18 months were collected and snap frozen in dry ice to perform the mRNA evaluation. mRNA was extracted using the RNAeasy Mini Kit (Qiagen) according to manufacturer’s protocol. The extraction was performed automatically using the QIAcube device from Qiagen. RNA concentration was subsequently quantified using a NanoDrop 2000C. Samples with a RIN value under 5 were excluded. cDNA synthesis was performed using Superscript Vilo cDNA Synthesis (ThermoScientific) according to the manufacturer’s protocol. TaqMan^®^ OpenArray^®^ Mouse Inflammation, TaqMan^®^ OpenArray^®^ Real-Time PCR Master and TaqMan^®^ OpenArray^®^ Real-Time PCR were used to performe the qPCR. Real-Time PCR Open Array from Applied Biosystems was used to read the Open Array 384 well plate used to perform the qPCR.

### Gene array analysis

Differently expressed genes are represented using the data from the HTqPCR assay assessed in the *Openarray* platform (Qiagen). The statistical analysis was performed using the software *DataAssist v3.01.* The maximum CT permitted was 35. First, we sorted the data based on the Gene Fold Change and then we convert the data to Log2FC. We compared the 5xFAD at 6 and 18 months to WT mice and 5xFAD with 5xFAD Gal3KO at 6 and 18 months as well. We selected genes with a Log2FC value between ±2 at 6 months and ±4 at 18 months. 95 genes out of 629 were selected at 6 months and 106 genes at 18 months. For the analysis of the main pathways affected by the lack of Gal3 at 6 and 18 months in our 5xFAD mouse model we used Network analysis along with KEGG and Reactome database.

### Behavioral tests

The Morris water maze test tests spatial acquisition memory and was conducted in a pool consisting of a circular tank (180 cm diameter) filled with opaque water at 20 ° C ± 1 °C) A platform (15 cm diameter) was submerged 10 mm under the water surface. A white curtain with specific distal visual cues surrounded the water maze. White noise was produced from a radio centrally positioned above the pool to avoid the use of auditory cues for navigation. Spatial learning sessions were conducted on the following 10 consecutive days with four trials per day. Each trial was started by introducing the mouse, facing the pool wall, at one of four starting points in a quasi-random fashion way to prevent strategy learning. Each mouse remained on the platform for 30 sec before transfer to a heated waiting cage. During all acquisition trails, the platform remained in the same position. On the day following the last learning trial, a 60 sec probe test was conducted, during which the platform was removed from the pool. All mouse movements were recorded using computerized tracking system that calculated distances moved and latencies required for reaching the platform (ANY-maze 5.2)

### Total RNA extraction and qPCR

Total RNA and proteins were extracted using TriPure Isolation Reagent (Roche). RNA integrity (RIN) was determined by RNA Nano 6000 (Agilent). The RIN was 8.5 ± 0.5. RNA was quantified using a NanoDrop 2000 spectrophotometer (Thermo Fischer, Spain).

#### Retrotranscription and quantitative real-time RT-PCR

Retrotranscription (RT) (4 μg of total RNA) was performed with the High-Capacity cDNA Archive Kit (Applied Biosystems). For real-time qPCR, 40 ng of cDNA were mixed with 2× Taqman Universal Master Mix (Applied Biosystems) and 20x Taqman Gene Expression assay probes (Applied Biosystems, supplemental). Quantitative PCR reactions (qPCR) were done using an ABI Prism 7900HT (Applied Biosystems). The cDNA levels were determined using GAPDH as housekeeper. Results were expressed using the comparative double-delta Ct method (2-ΔΔCt). ΔCt values represent GAPDH normalized expression levels. ΔΔCt was calculated using 6-month-old WT mice samples.

#### Primers

Iba1 (Ref. Mm00479862_g1), CD45 (Ref. Mm01293577_m1), CD68 (Ref. Mm03047343_m1), TREM2 (Ref. Mm04209424_g1), Cx3Cr1 (Ref. Mm02620111_s1), GAPDH (Ref. Mm99999915_g1), IL-6 (Ref. Mm00446190_m1), TNFa (Ref. Mm00443258_m1), GFAP (Ref. Mm01253033_m1).

### Genetic association analysis

#### Datasets

Genotypic datasets from four GWAS were used in this study: a) The Murcia study [2]; b) The Alzheimer’s Disease Neuroimaging Initiative (ADNI) study [49]; c) The GenADA study [44]; and d) The NIA study [69]. (For GWAS dataset details see supplementary information.) The Murcia study was previously performed by researchers from our group [3]. Datasets from ADNI, GenADA, and NIA, studies were obtained from dbGAP (http://www.ncbi.nlm.nih.gov/gap), Coriell Biorepositories (http://www.coriell.org/) or ADNI (http://adni.loni.ucla.edu/). Prior to the genetic association analysis, each dataset (Murcia, ADNI, GenADA, NIA, and TGEN) was subjected to both an extensive quality control analysis and a principal component analysis. In addition, since different platforms were used in the five GWAS analyzed, we imputed genotypes using HapMap phase 2 CEU founders (n = 60) as the reference panel. These approaches have been previously described [3, 9, 46]. Overall, a total of 2252 cases and 2538 controls were included in the meta-analysis.

#### SNP selection

To select single nucleotide polymorphisms (SNPs) within *LGALS3* gene including 1000 pb upstream and downstream of that genetic region we used the UCSC Table Browser data retrieval tool[34], release genome assembly: Mar. 2006 (NCBI36/hg18), from the UCSC Genome Browser database (http://genome.ucsc.edu/)[29]. Selected SNPs were extracted from GWAS datasets using Plink v1.06 software[56].

#### Linkage disequilibrium blocks

Linkage disequilibrium (LD) blocks were determined along the genomic regions studied using Haploview software [6] and genotyping data from the largest dataset used (NIA dataset).

#### Association analyses

Unadjusted single-locus allelic (1 df) association analysis within each independent GWAS dataset was carried out using Plink software. We combined data from these four GWAS datasets using the meta-analysis tool in Plink selecting only those markers common to, at least, three studies. For all, single locus meta-analyses, fixed effects models were employed when no evidence of heterogeneity was found. Otherwise random effects models were employed.

All selected SNPs were located close to 3’ end of *LGALS3* gene. Because all of them belonged to the same linkage disequilibrium block, multiple testing correction was not applied. Thus, the p-value threshold was established 0.05.

### Supplementary information for GWAS study

Description of genome-wide association study (GWAS) datasets employed in the study:

The Murcia study was designed as a new case-control GWAS in the Spanish population. In this study, 1,128 individuals were genotyped using Affymetrix NspI 250K chip. A sample of 327 sporadic Alzheimer’s disease (AD) patients diagnosed as possible or probable AD in accordance with NINCDS-ADRDA criteria by neurologists at the Virgen de Arrixaca University Hospital in Murcia (Spain) and 801 controls with unknown cognitive status from the Spanish general population were included. The Alzheimer’s Disease Neuroimaging Initiative (ADNI) longitudinal study was launched in 2003 by the National Institute on Aging (NIA), the National Institute of Biomedical Imaging and Bioengineering (NIBIB), the Food and Drug Administration (FDA), private pharmaceutical companies, and non-profit organizations, as a $60 million, 5-year public-private partnership (1). The primary goal of ADNI has been to test whether serial magnetic resonance imaging, positron emission tomography, other biological markers, and clinical and neuropsychological assessment can be combined to measure the progression of mild cognitive impairment (MCI) and early AD. Determination of sensitive and specific markers of very early AD progression is intended to aid researchers and clinicians to develop new treatments and monitor their effectiveness, as well as lessen the time and cost of clinical trials. The Principal Investigator of this initiative is Michael W. Weiner, MD, VA Medical Center and University of California – San Francisco. ADNI is the result of efforts of many co-investigators from a broad range of academic institutions and private corporations, and subjects have been recruited from over 50 sites across the U.S. and Canada. The initial goal of ADNI was to recruit 800 adults, ages 55 to 90, to participate in the research, approximately 200 cognitively normal older individuals to be followed for 3 years, 400 people with MCI to be followed for 3 years and 200 people with early AD to be followed for 2 years. For up-to-date information, see www.adni-info.org. The GenADA study included 801 cases meeting the NINCDS-ADRDA and DSM-IV criteria for probable AD, and776 control subjects without family history of dementia that were genotyped using Affymetrix 500K GeneChip Array set. The NIA Genetic Consortium for Late Onset Alzheimer’s Disease (LOAD) Study originally included 1,985 cases and 2,059 controls genotyped with Illumina Human 610Quad platform. Using family trees provided in the study, we excluded all related controls and kept one case per family. A total of 1,077 cases and 876 controls.

### Human material

All the human material used has been obtained from the Lund University Hospital, Neuropathology Unit (Table 1) (Elisabet Englund) and The Netherlands Institute for Neuroscience, Amsterdam, The Netherlands (Inge Huitinga) (Table 2)

Written informed consent for the use of brain tissue and clinical data for research purposes was obtained from all patients or their next of kin in accordance with the International Declaration of Helsinki. Medisch Ethische Toetsingscommissie (METc) of VU University has approved the procedures of brain tissue collection and the regional ethical review board in Lund has approved the study. All human data was analyzed anonymously.

### Antibodies

Antibodies used for this study: Anti-rabbit iNOS primary Antibody (1:5000, Santa Cruz), Anti-rat Gal3 Antibody, (1:3000, M38 clone from Hakon Leffler’s lab, (in-house antibody) (Add references), Anti-Goat Gal3 Antibody (1:1000, R&D Systems) Anti-mouse Actin antibody 1:10000 (Sigma-Aldrich), Anti-human Aβ antibody (1:5000, Covance), Anti-Rabbit Iba-1 antibody (1:500, Wako), Anti-mouse TLR4 Antibody (1:1000, Santa Cruz), Anti-mouse NLRP3 Antibody (1:5000, Adipogen), Anti-rabbit C83 antibody (369) (1:1000, Gunnar Gouras Laboratory, BMC, Lund, Sweden)(Add References) Anti-rabbit IDE-1 antibody (1:1000, Calbiochem), Anti-Rabbit p-Tau (pTau181, Santa Cruz), Anti-mouse Aβ (1:1000, Sigma-Aldrich). Anti-CD45 rat monoclonal (clon IBL-3/16, 1:500 dilution, AbD Serotec). Secondary antibodies used for western blot were: anti-rabbit, anti-mouse, anti-goat and anti-rat from Vector Labs. Secondary antibodies used for immunofluorescence raised in donkey were: anti-rabbit, anti-goat, anti-mouse and anti-rat from Life Technology (AlexaFluor).

### Inhibitor used in our study

The inhibitor used for the experiments in Fig. 2A-C and Suppl. Fig 5 was 1,1’-sulfanediyl-bis-{3-deoxy-3-[4-(3-fluorophenyl)-1*H*-1,2,3-triazol-1-yl]-β-D-galactopyranoside} [54] (inhibitor 1), and it was synthesized and characterized as reported previously [17]. The purity was determined to be 97.3% according to UPLC-analysis (Waters Acquity UPLC system, column Waters Acquity CSH C18, 0.5 ml/min H_2_O-MeCN gradient 5-95% 10 min with 0.1% formic acid).

## RESULTS

### Galectin-3 is expressed in microglial cells positioned next to amyloid plaques in AD human cortical samples and 5xFAD mice

We first evaluated the levels of gal3 in cortical sections from AD patients and age-matched healthy controls (Fig. 1A). We found a 10-fold increase of gal3 in AD patients compared to controls. Next, we performed gal3, Aβ and microglial (Iba1) staining on AD and healthy cortical sections (Fig. 1B-C). The expression of gal3 was mostly absent in the control samples with a faint staining in brain blood vessels (Fig. 1C). However, in the AD cortical samples, there was a robust gal3 staining in Iba1-positive cells (Fig. 1C), which was strictly confined to Aβ plaque-associated microglia (Fig. 1B). To further confirm our observations, we used 5xFAD, a transgenic AD mouse model and found the expression of gal3 significantly upregulated in a time-dependent fashion from 6 to 18 months (Fig. 1D; Supplementary Fig. 1). In the 5xFAD brains, gal3 was typically found in microglial cells associated to Aβ plaques (Fig. 1E) Other cell types, such as neurons and astrocytes, very rarely expressed gal3 (Suppl. Fig. 2A-C).

**Fig. 1.**
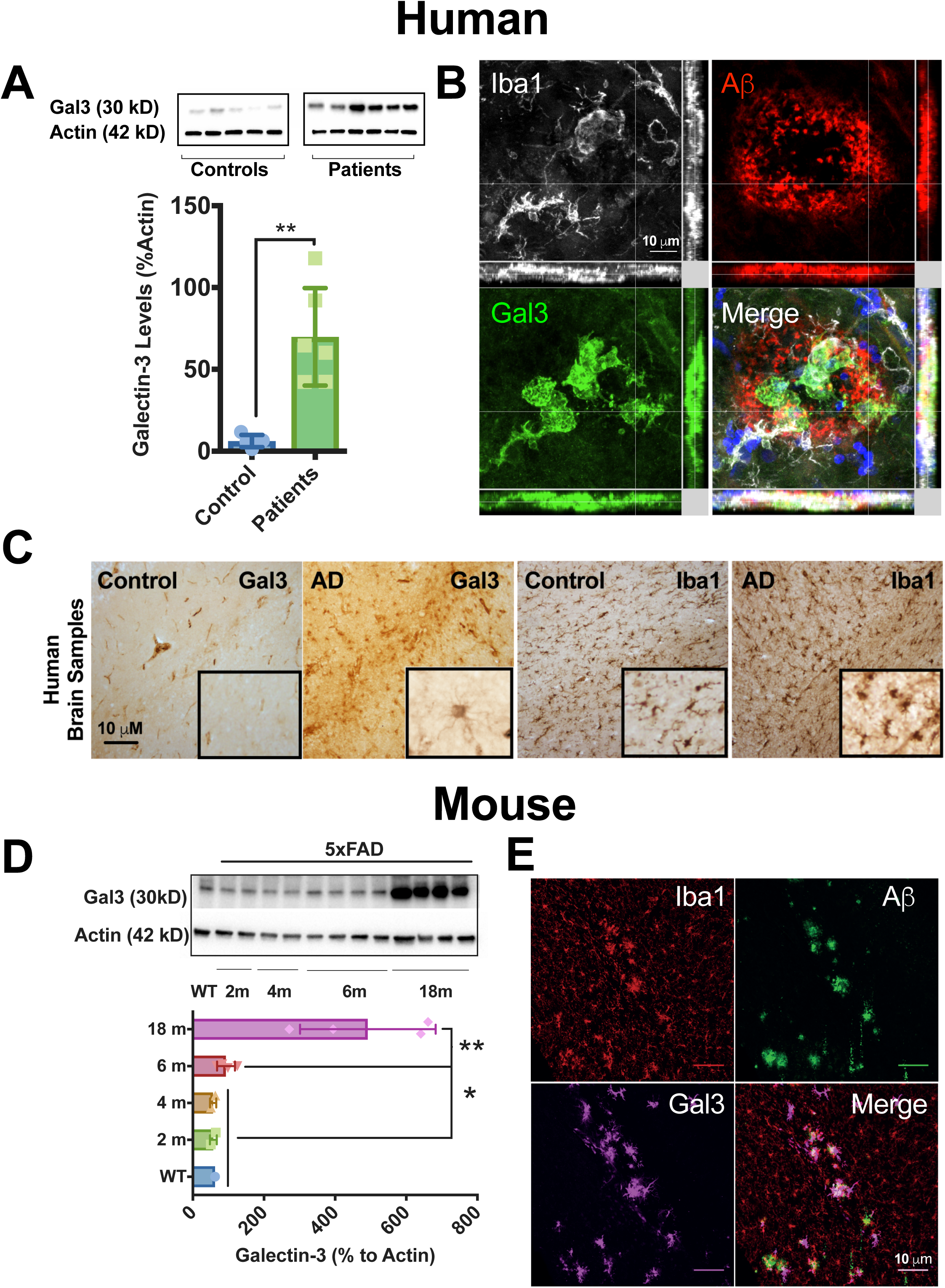

### Single nucleotide polymorphisms (SNP) associated with the gal3 gene increase the risk of developing AD

We performed genetic association studies using a SNP meta-analysis approach of the *LGALS3* gene, which encodes gal3. A total of 60 SNPs were identified in the *LGALS3* genetic region. Only five SNPs were genotyped in at least three GWAS (Suppl. Fig. 3A, Table 3), being in high linkage disequilibrium (d’ > 0.96; Suppl. Fig. 3B). Interestingly, all five SNPs were associated with an increased AD frequency (p < 0.03; Table 3), suggesting a potential causal role of gal3 in the development of AD.

### Galectin-3 regulates amyloid-dependent microglial activation

To investigate the role of gal3 in microglial activation in AD, we first evaluated the inflammatory response in BV2 microglial cells. Aβ42 fibrils were characterized prior to the experiments (Suppl. Fig 5A). Our preparation was tested for LPS endotoxin as well as for cell toxicity (Suppl. Fig. 5B-C). BV2 cells challenged with Aβ42 fibrils activated microglial cells, inducing the expression of iNOS and NLRP3 (Suppl. Fig. 5D) and the production and release of proinflammatory cytokines (Suppl. Fig. 5E). Next, we challenged BV2 cells with Aβ42 fibrils at 10 μM along with increasing concentration of gal3 inhibitor (10 and 25 μM) for 12 h. The conditioned medium was collected and inflammatory-related cytokines were measured. Notably, TNFα, IL12, IL6 and IL8 proinflammatory cytokine release was significantly reduced in response to gal3 inhibition in a concentration-dependent manner (Fig. 2A). The same was true for iNOS, a classical proinflammatory marker as measured by western blot (Fig. 2B). We also evaluated the levels of Insulin Degrading Enzyme 1 (IDE-1) on BV2 cells. IDE-1 is an enzyme involved in Aβ degradation [57]. IDE-1 was downregulated in BV2 cells challenged with Aβ42 fibrils in a concentration-dependent fashion (Fig. 2C, left). Strikingly, with gal3 inhibition, the downregulation of Aβ-induced IDE-1 turned into a significant upregulation in a concentration-dependent manner (Fig. 2C, right). To further confirm our *in vitro* data, we evaluated the inflammatory response using primary microglial cultures from WT and Gal3KO mice. Primary microglial cultures were challenged with Aβ42 fibrils at 3 and 10 μM. Similar to BV2 cells, the lack of gal3 reduced the release of proinflammatory cytokines such as IL6, IL8 and TNFα (Fig. 2D). Other cytokines such as IFNγ, IL4 and IL12 were not affected by the lack of gal3. We next analyzed the effect of gal3 in the phagocytosis Aβ fibrils by primary microglia. To achieve this, primary microglial cells were first pre-treated (30 min) with gal3 (1 μM) and then challenged with Aβ fibrils. Microglial uptake of Aβ fibrils was reduced in gal3-treated primary cultures (Fig. 2E).

**Fig. 2.**
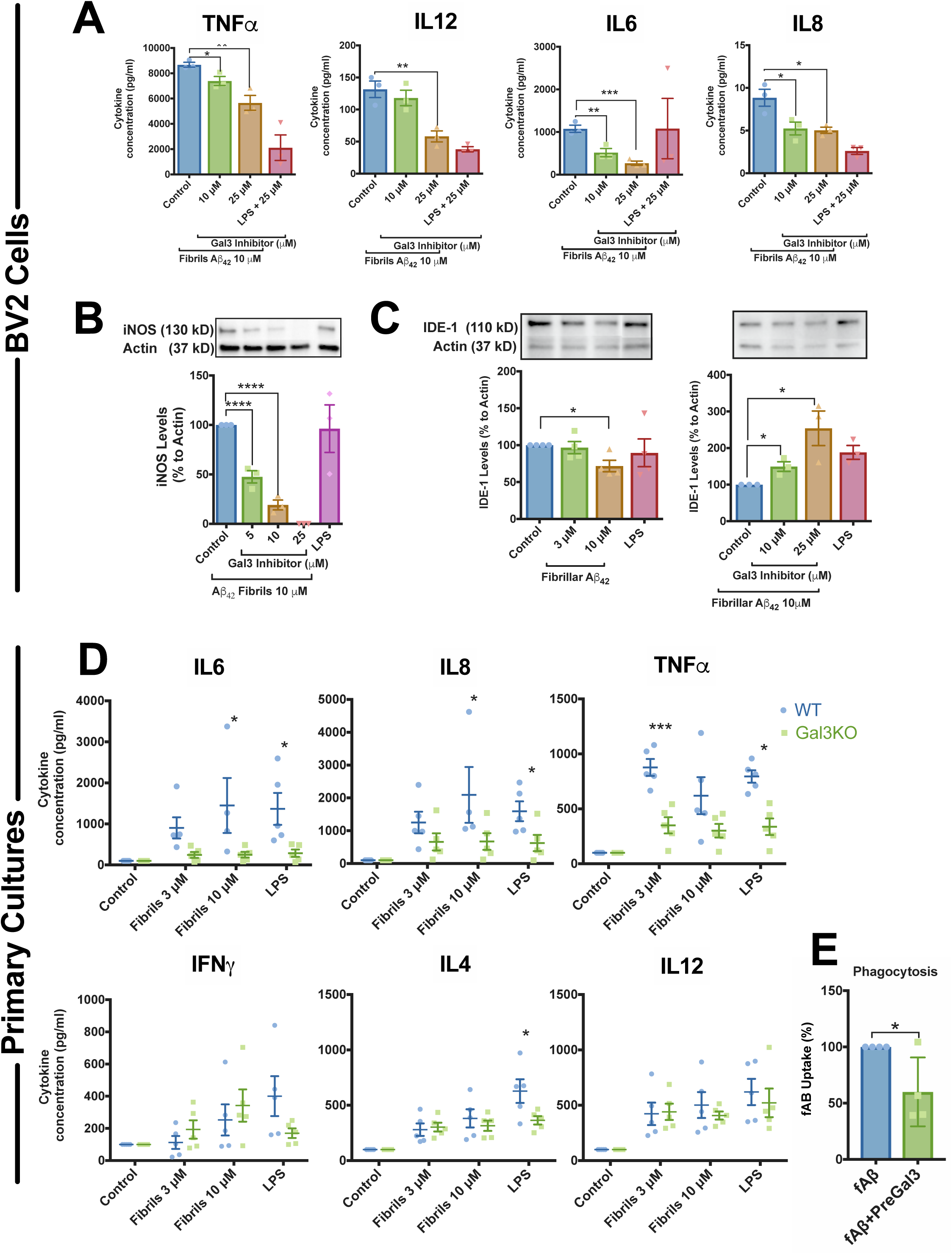

### Galectin-3 regulates microglial activation in 5xFAD mice

To examine the role of gal3 in the neuroinflammatory response, WT, 5xFAD mice and crossbred 5xFAD/Gal3KO mice were compared. Transcriptional profiles of hippocampal tissue were obtained by running high-throughput qPCR against a 632-gene mouse inflammation panel. WT and gal3KO mice showed quite similar profiles, suggesting a limited role of gal3 in promoting inflammation in the absence of pathology. In 5xFAD mice, compared to WT, a significant number of genes were affected, especially in 18-month-old mice, in which 125 out of 629 immune genes were highly upregulated. These included complement factor, chemokines, interleukin receptors and toll-like receptors (Suppl. Fig. 4). Overall, the whole immune response was highly attenuated in 5xFAD/Gal3KO mice (Suppl. Fig. 4). Interestingly, genes associated with the recent characterized neurodegenerative disease-associated phenotype (DAM microglia[35, 37]) such as Clec7a, Csf1, Cd74, Cxcl10 and Cybb (Fig. 3A, B and D) were downregulated in 5xFAD/Gal3KO compared to 5xFAD. qPCR analysis of homeostatic (Cx3cr1, TGFbeta1) and DAM/reactive microglia (CD45, TREM2, Clec7a, CD68, TNFα and Lysozyme M) confirmed the instrumental role of gal3 in driving the brain immune response in 5xFAD mice (Fig. 3). To identify potential cell signaling pathways related to gal3 in 5xFAD mice, we used NetworkAnalyst (http://www.networkanalyst.ca)[70]. Most affected pathways were related to TLR and DAP12 signaling (Fig. 3B and D). DAP12 has been described as a TREM2 adaptor, which plays a critical role in the switch from homeostatic to disease-associated [35, 37]. Our data anticipates a key role of gal3 in driving microglia activation in AD through regulation of different microglial pattern-recognition receptors, including TLRs and TREM2.

**Fig. 3.**
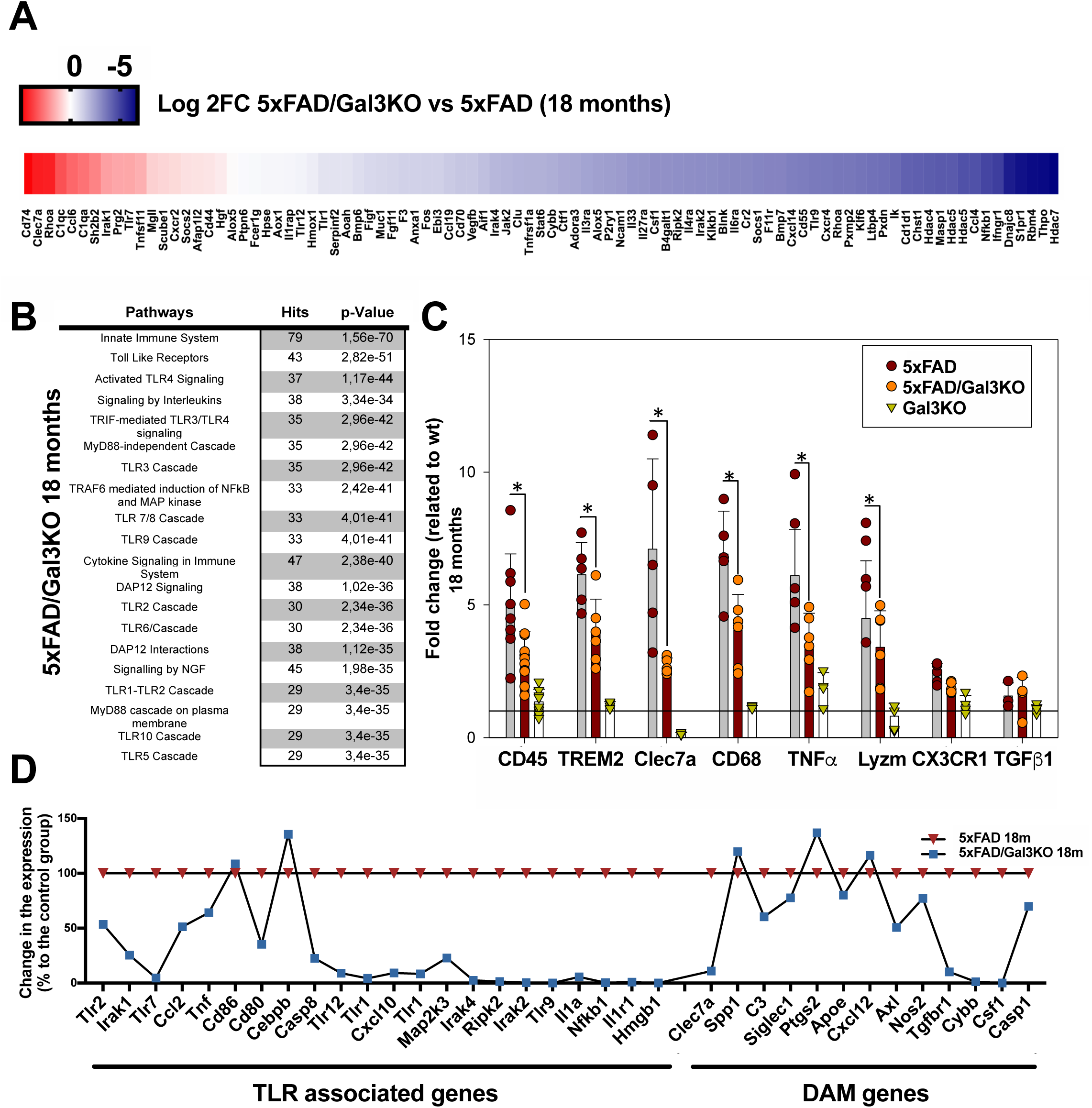

### The lack of galectin-3 reduces AD pathology in 5xFAD mice

Once we confirmed the important role of gal3 in driving AD-associated microglial immune responses, we next analyzed the effect of gal3 in AD pathogenesis. There was clear age-dependent Aβ deposition in 5xFAD mice (Fig. 4). Aβ deposition in aged animals (18 months) was very high, an indication that Aβ clearance becomes saturated in aged 5xFAD mice. Consequently, in order to detect a potential effect of gal3 in Aβ burden, we analyzed the effect of gal3 gene deletion in two well-defined areas, hippocampal CA1 area (high rate of Aβ deposition) and thalamus (low rate of Aβ deposition), in 5xFAD mice at 6 and 18 months. Notably, the plaque burden was significantly reduced in 5xFAD mice lacking gal3 at 6 months at CA1 (Fig. 4A). At 18 months, we could not find any difference between both groups, probably due to oversaturated plaque deposits in the hippocampi (Fig. 4B). In agreement with this view, a significant reduction in Aβ plaque deposition was found in the thalamus in 5xFAD/Gal3KO mice at 6 and 18 months compared to aged-matched 5xFAD controls (Fig. 4A-B).

**Fig. 4.**
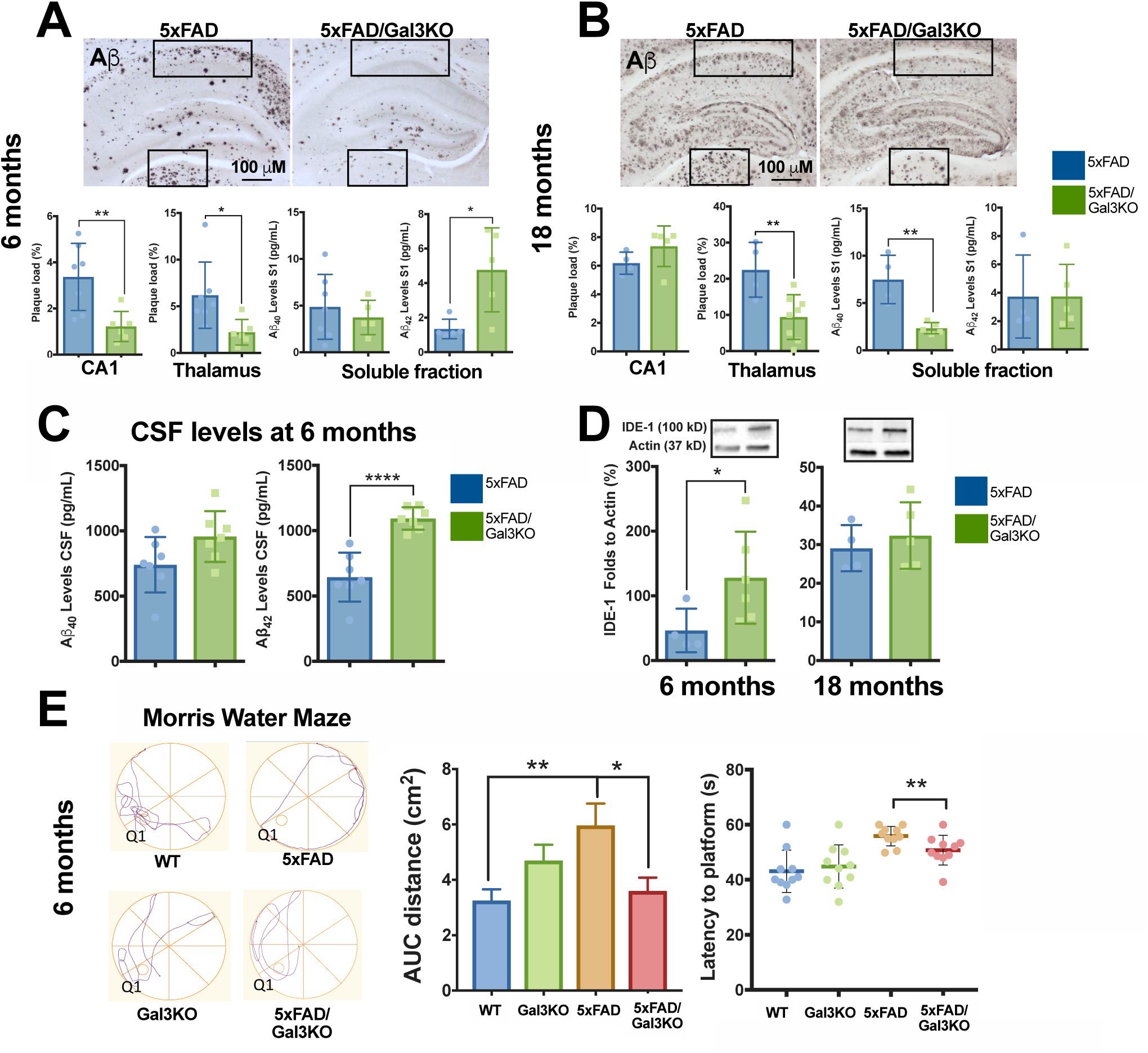

Next, we measured cortical Aβ42 and Aβ40 levels from soluble brain fractions. The levels of Aβ40 were significantly lower in 5xFAD/Gal3KO (vs. 5xFAD) brains at 18 months but not at 6 months (Fig. 4A-B). Conversely, Aβ42 levels were significantly higher in 5xFAD/Gal3KO (vs. 5xFAD) brains at 6 months (Fig. 4A) with no change at 18 months (Fig. 4B). In the insoluble Aβ fraction, we found reduced Aβ40 in 5xFAD/Gal3KO at 6 months (Suppl. Fig. 6A). Changes in Aβ40 and Aβ42 levels were not due to a change of APP expression as 5xFAD and 5xFAD/Gal3KO mice showed comparable levels at both ages measured (Suppl. Fig. 6B). Aβ is cleared from the brain by different mechanisms, including microglia phagocytosis, drainage to the ventricular system and activity of extracellular proteases, such as IDE-1 [26]. We found a significant increase of Aβ42 CSF levels in 5xFAD/Gal3KO mice compared to 5xFAD mice (Fig. 4C), mimicking the clinical situation in which Aβ in the CSF is inversely related to the Aβ plaque load in AD patients. Interestingly, hippocampal IDE-1 levels were increased in 5xFAD/Gal3KO at 6 months compared to 5xFAD (Fig. 4D), supporting our *in vitro* data (Fig. 2C).

Having established that gal3 is a main driver of the AD-associated inflammatory response and that gal3 deficiency reduces Aβ plaque burden, we next studied the impact of gal3 deficiency on cognitive behavior. The cognitive ability was evaluated using the Morris water maze memory test. Remarkably, the lack of gal3 in 5xFAD/Gal3KO mice resulted in better cognitive performance compared to 5xFAD mice as the 5xFAD/Gal3KO mice were able to navigate and reach the platform by using a shorter path (Fig. 4E).

### Galectin-3 drives microglia reactivity in Aβ plaques

In view of our findings that gal3-reactive microglia are strictly confined to Aβ plaques and that gal3 deficiency impairs the inflammatory response and Aβ plaque load, we next analyzed microglial coverage on the surface of amyloid plaques in cortex from both 5xFAD and 5xFAD/Gal3KO mice. First, we found that 5xFAD/Gal3KO mice showed reduced Iba1 immunoreactivity in the cortex compared to 5xFAD mice (Fig. 5A) with these differences being particularly evident in plaque-associated microglia in 5xFAD/Gal3KO mice (Fig. 5B). Remarkably, gal3 deficiency significantly reduced Iba1-labeled microglial clusters around Aβ plaques (Fig. 5C). Analysis of microglia coverage over plaque surface showed that a high subset of plaque-associated microglia (<50%) expressed high levels of gal3 (Fig. 5D). Interestingly, these findings are very similar to those seen in 5xFAD/TREM2KO mice [42, 48, 74]), in which TREM2 deficiency reduced the capacity of microglia to cluster around Aβ plaques. This data suggests that gal3-associated biological effects may be associated with TREM2 signaling.

**Fig. 5.**
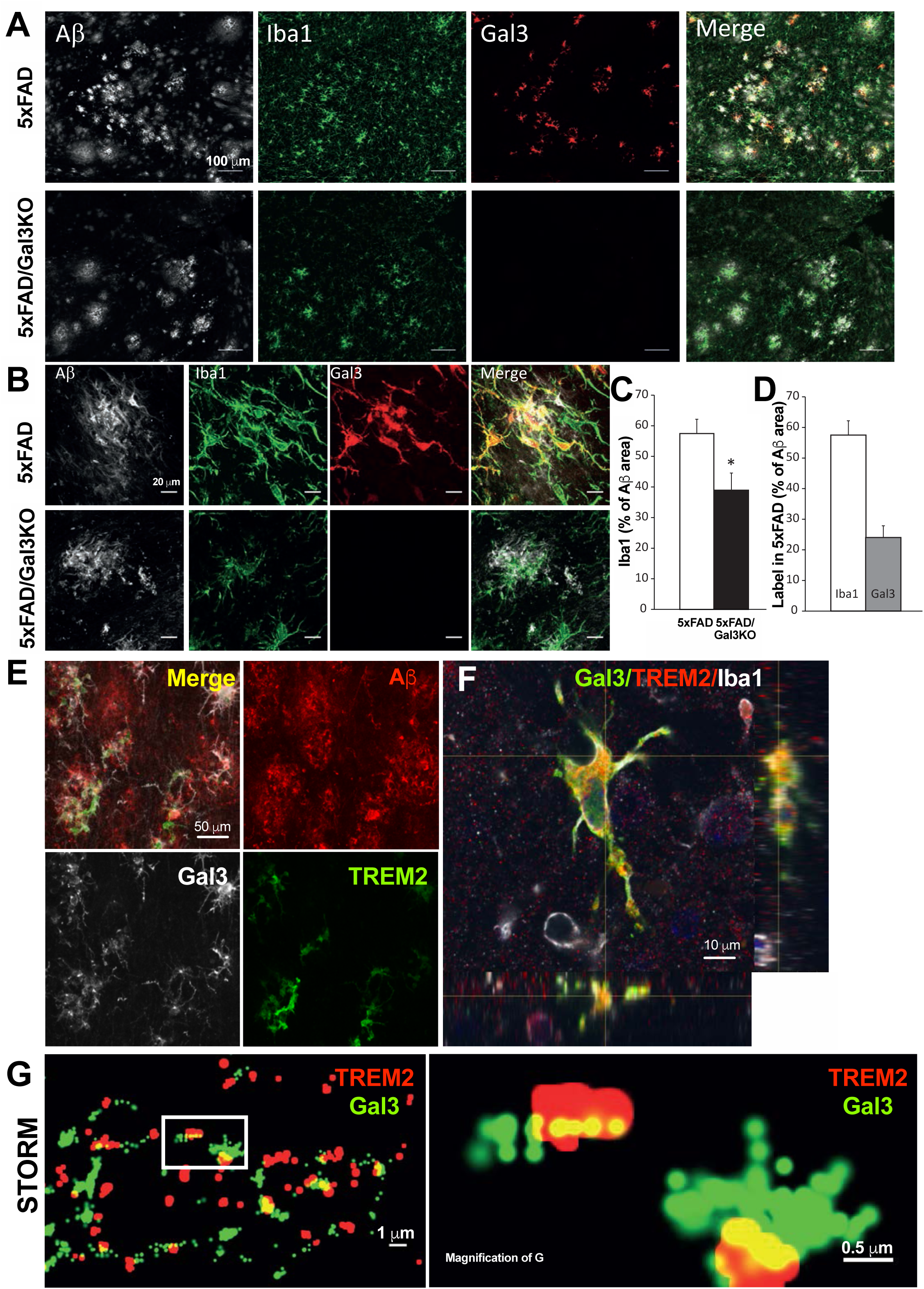

### Galectin-3 acts a TREM2 ligand

Recent single cell transcriptomic analyses of microglia under different conditions of neurodegeneration, including AD, have revealed gal3 as one of the most upregulated genes in processes that have been suggested to be TREM2-dependent[37, 47]. We have demonstrated that gal3 is released by reactive microglial cells[10] and may thus interact with different receptors controlling key functions of microglial cells, including TLR4[47] and MerTK[53]. One of the main pathways altered in 5xFAD/Gal3KO mice was related to DAP12 signaling (Fig. 3B). Since DAP12 is a downstream regulator of TREM2, we wondered if gal3 could act as an endogenous TREM2 ligand. To test our hypothesis, we first performed TREM2, gal3 and 6E10 staining on 5xFAD brain sections and found i) the majority of TREM2-labelled microglia was gal3+ (83.4 ± 5.7% and 94.0 ± 3.02% at 6 and 18 months, respectively), an indication that gal3 specifically labels the DAM phenotype [80] (Fig. 5E, Suppl. Fig. 7A), and ii) a striking cellular co-localization of TREM2 and gal3 in microglial cells around Aβ plaques (Fig. 5E-F, Suppl. Fig. 7A). We next used quantitative high-resolution confocal microscopy and super-resolution Stochastic Optical Reconstruction Microscopy (STORM) to determine whether TREM2 and gal3 physically interact using brain sections from 5xFAD mice. Our ultrastructural analysis clearly revealed the physical interaction between gal3 (green) and TREM2 (red) over the membrane surface of microglia (see yellow dots in Fig. 5G). Further, we incubated brain sections from 5xFAD/Gal3KO with recombinant gal3 to further analyze gal3 binding in situ. This experiment demonstrated the binding of gal3 to TREM2^+^ but not in TREM2^-^ /Iba1-labelled microglial cells (Suppl. Fig. 7B).

We further assessed the direct TREM2-gal3 interaction by the potency of soluble TREM2 to inhibit binding of a fluorescent glycosylated probe to gal3 as measured by fluorescence anisotropy. This analysis demonstrated a strong interaction between WT gal3 and TREM2 with a Kd value of 450 ± 90 nM (Fig. 6A) in the same range as other preferred glycoproteins [12, 13]. A mutant gal3 (R186S, deficient in the carbohydrate binding domain, with severely reduced affinity for LacNAc-structures) [58] bound TREM2 much more weakly (Kd of 5900 ± 600 nM; Fig. 6A). This means that gal3 interacts strongly with TREM2 protein [Kd in the same range as *e.g.* some serum glycoproteins such as transferrin and haptoglobin[12, 13] and that the interaction is carbohydrate dependent. The hypothetic interaction was modeled based on previously reported TREM2 and gal3 structures (Suppl. Fig. 7C). Next, we tested whether gal3 could affect TREM2 signaling using a TREM2-DAP12 reporter cell line, and we found that added gal3 triggered TREM2-DAP12-dependent signaling in a dose-dependent manner (Fig. 6B). Overall, our data demonstrate that gal3 is an endogenous ligand for TREM2.

**Fig. 6.**
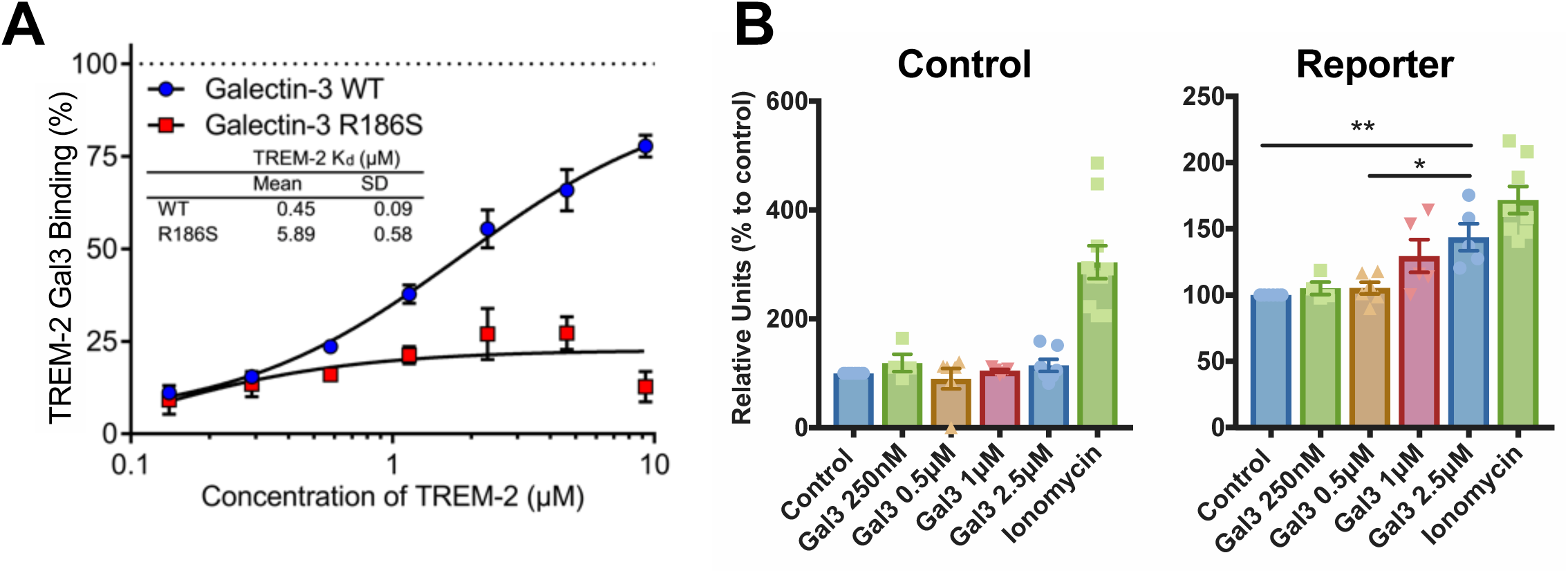

### Galectin-3 acts as a cross-seeding agent of Aβ aggregation at nanomolar range

We have previously demonstrated that gal3 can be released by reactive microglial cells *in vitro* [10, 53]. However, to our knowledge, evidence supporting this view *in vivo* is lacking. To address this, we performed electron microscopy immunogold labeling of gal3 in sections from 12-month-old APP/PS1 mice. This ultrastructural study first demonstrated that immunogold particles specifically labelled microglial cells in the vicinity of Aβ plaques (Fig. 7A). Remarkably, immunoelectron microscopy confirmed gal3-labeling in the extracellular space and in Aβ fibrillar aggregates (Fig. 7B-C). The presence of extracellular gal3 associated to Aβ fibrillar aggregates is similar to that seen for ASC specks (NLRP3 inflammasome platform), which are released by plaque-associated microglia and induce Aβ cross-seeding[66]. Intriguingly, TREM2 has been associated with Aβ plaque compaction[74], and, hence, the possibility emerged that gal3 may have a role in Aβ aggregation. We, thus, tested whether gal3 affects aggregation of pure Aβ monomers using a thioflavin T assay, in which thioflavin T fluoresces when bound to beta-sheet-containing aggregates. We used a physiological medium and conditions in which 10 μM Aβ aggregated very little over 15 hours, but the addition 1 nM of gal3 increased aggregation as measured by bulk fluorescence (Fig. 7D) or microscopy (Fig. 7E and F). The increased thioflavin T fluorescence could be removed by centrifugation (Fig. 7G), indicating that it was due to the presence of aggregates. Note, that this increased aggregation was at a molar ratio of gal3 to Aβ of 1:10 000, suggesting that gal3 increased nucleation of Aβ aggregation.

**Fig. 7.**
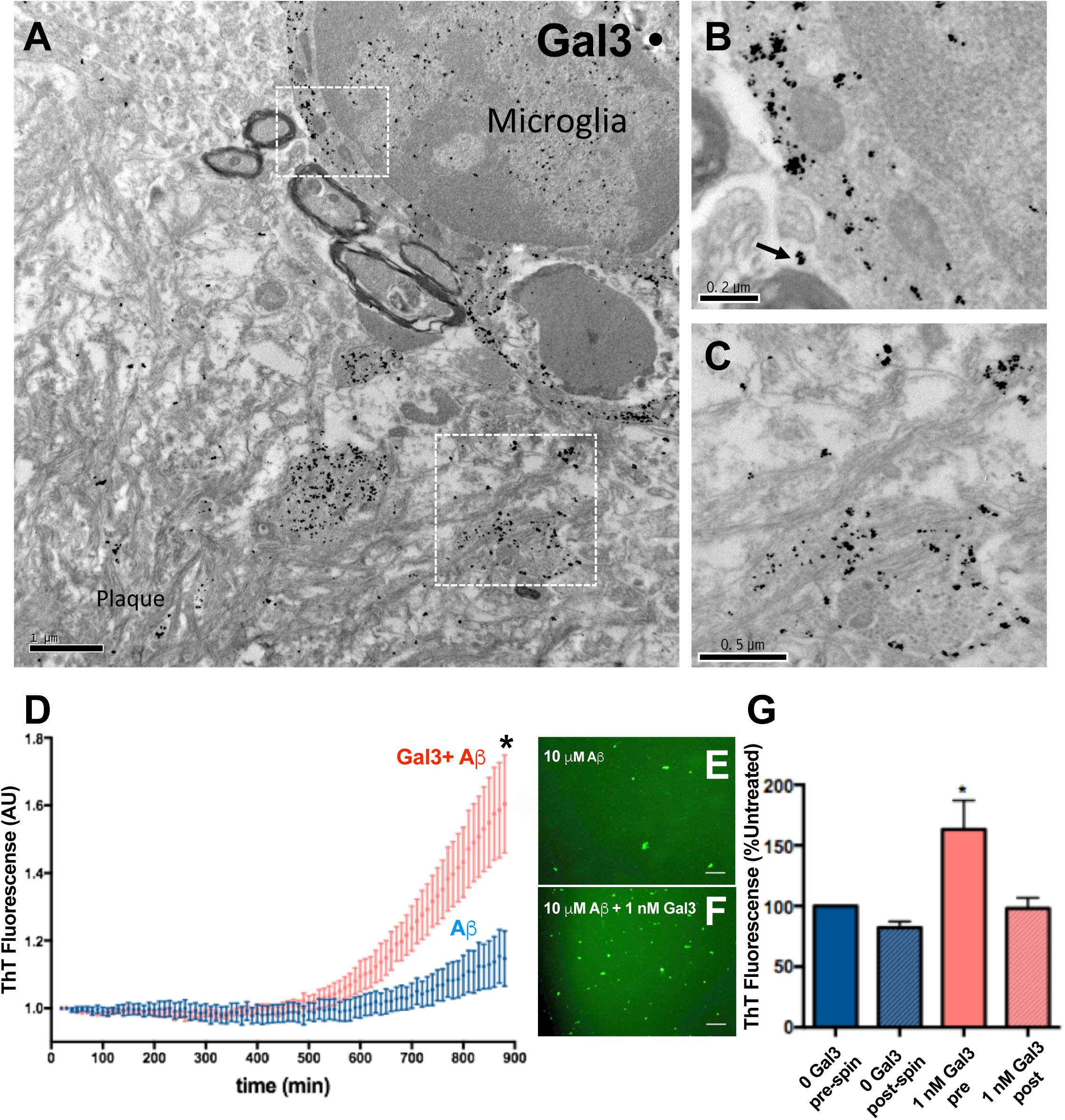

### Galectin-3 induces the formation of insoluble Aβ aggregates following injections of Aβ monomers in the hippocampi of WT mice

To evaluate if gal3 acts as a cross-seeding agent of Aβ *in vivo,* Aβ42 monomers (10 μM) were incubated alone or in the presence of Gal3 (10 μM) for 1 hour at 37 °C. Then, 2 μls were injected in either the left (Aβ+gal3) or the right hippocampi (Aβ alone) of WT mice. The analysis was performed two months post-injection. The injection of Aβ monomers alone failed to show any sign of Aβ aggregation in the injected hippocampi (Fig. 8A). In sharp contrast and remarkably, the presence of Aβ-like plaques were evident in the hippocampi injected with Aβ+gal3 (Fig. 8B). These Aβ aggregates were positive for gal3, Aβ (6E10 antibody) and thioflavin-S (Fig. 8E), thus demonstrating the amyloid structure of the Aβ deposits. These amyloid deposits were surrounded by Iba1-labelled microglia (Fig. 8F) and GFAP-positive reactive astrocytes (Fig. 8G). Our data unequivocally demonstrates that gal3 is a microglial-associated cross-seeding agent of Aβ aggregation.

**Fig. 8.**
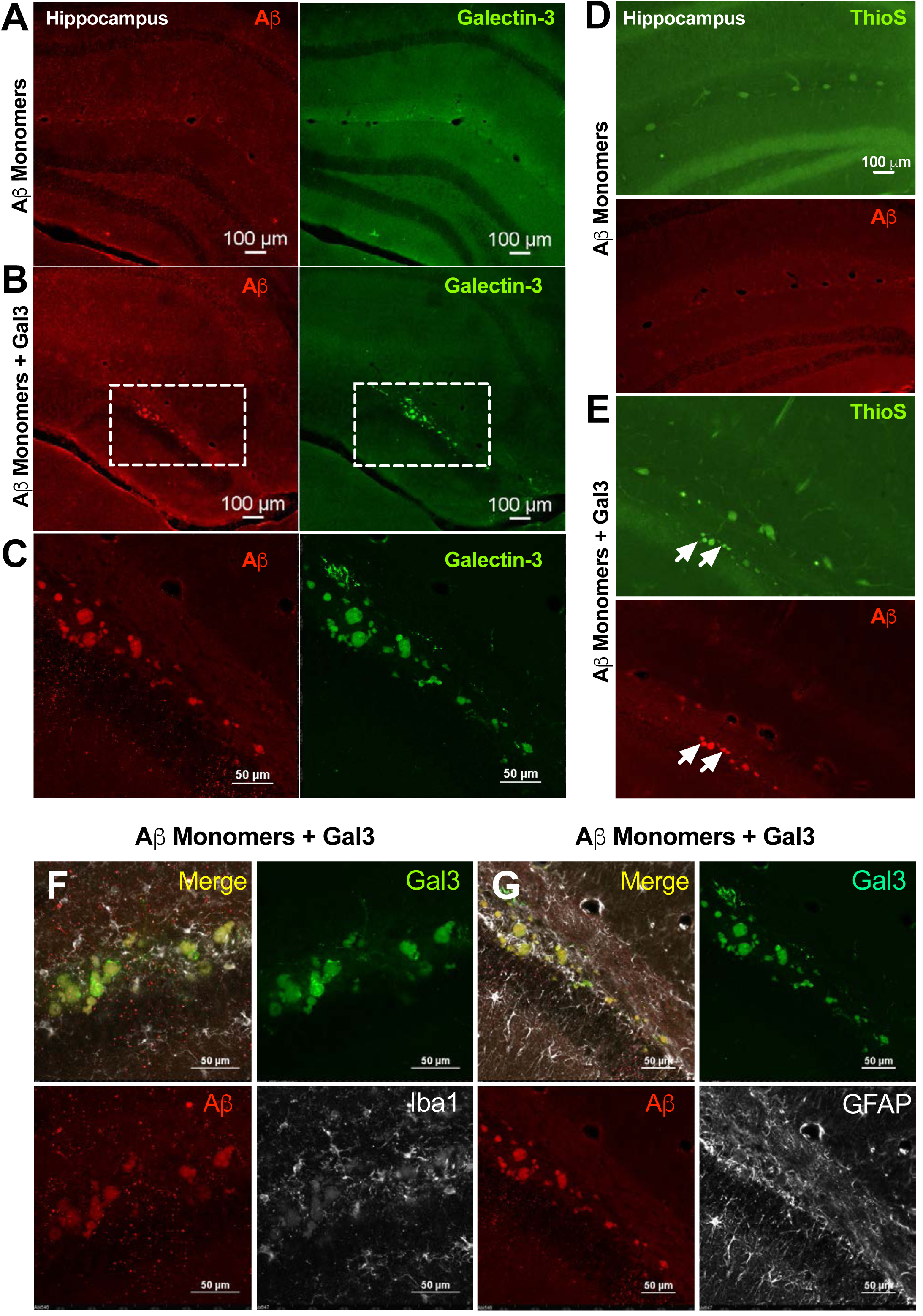

## DISCUSSION

Recent findings suggest an important role of neuroinflammation in neurodegenerative diseases [25]. Thus, selective mutations in microglia/myeloid-specific genes, including the glycoprotein triggering receptor on myeloid cells 2 (TREM2), have been associated with AD [22]. Experimental AD studies have suggested that TREM2 is instrumental in neuroinflammation [32, 43] and drives the recently identified disease-associated microglia (DAM phenotype) [35, 37]. Gal3, a carbohydrate-binding protein, is one of the most upregulated genes associated with DAM [37, 42, 47]. Holtman et al. (2015) anticipated gal3 as an instrumental gene in driving the DAM program [68]. In order to study the role of gal3 in AD pathogenesis, we have analyzed human brains from AD patients and 5xFAD mice lacking gal3. The main conclusions from our study are i) gal3 protein is increased 10-fold in human AD brains and is mostly restricted to plaque-associated microglia in humans and 5xFAD mice, ii) SNPs associated with the gal3 gene are increased in AD patients, iii) gal3 deficiency reduces Aβ plaque burden and overall proinflammatory response and improves cognitive performance in 5xFAD/Gal3KO mice, iv) gal3 is an endogenous TREM2 ligand and v) gal3 acts as a cross-seeding agent of Aβ *in vitro* and *in vivo.* Hence, gal3 emerges as a central upstream regulator of AD-associated pathology.

Given the relationship between innate immune-related genes and AD incidence [26, 27], deciphering how Aβ aggregates triggers neuroinflammation is of critical importance. For this purpose, we first analyzed the role of gal3 in Aβ-induced inflammatory response in microglial cells. To this end, we took advantage of both chemical inhibition and gene deletion. Gal3 inhibition robustly reduced fAβ-induced iNOS expression in BV2 cells, an effect that was extended to proinflammatory cytokines, including TNFα and IL6, IL8 and IL12, which we confirmed in primary microglia cultures from WT and Gal3 KO mice. This data suggests that gal3 is a critical alarmin that amplifies the Aβ-induced proinflammatory response. Since inefficient clearance of Aβ may play a determinant role in AD pathogenesis [76], we analyzed the effect of gal3 in two major mechanisms associated to Aβ clearance: fAβ phagocytosis and IDE-1 levels, a key metalloprotease involved in Aβ degradation by microglia [25, 28, 39]. Gal3 was able to reduce Aβ phagocytosis, and gal3 deficiency increased IDE-1 levels, suggesting a detrimental role of gal3 in Aβ clearance. Overall, our *in vitro* study demonstrates an instrumental role of gal3 in driving the fAβ-induced pro-inflammatory response and a deleterious effect in Aβ clearance.

Recent transcriptomic analysis of microglia at the single cell level has identified a common disease-associated microglia phenotype, which has been suggested to be driven by TREM2 [35, 37, 47]. Interestingly, Krasemann et al. (2017) identified gal3 as one of the most upregulated genes in plaque-associated microglia, supporting our findings in human AD brains and 5xFAD mice [37]. Recently, Yang and colleagues overexpressed TREM2 in 5xFAD mice (5xFAD/TREM2) and found gal3 as one of the main genes affected by TREM2 overexpression [41], suggesting that gal3 plays an important role in microglia function under disease conditions. We generated 5xFAD/Gal3KO mice to answer whether gal3 plays a role in microglia-associated AD pathogenesis and to test if gal3 signaling is associated with TREM2. To answer this question, we performed an inflammatory gene array, demonstrating an age-dependent inflammatory response, including complement components, chemokines, interleukin receptors, toll-like receptors and DAM genes in 5xFAD mice. This inflammatory response was highly attenuated in 5xFAD/Gal3KO mice, thus confirming gal3 as a master regulator of AD-associated brain immune responses. Pathway analysis at 6 and 18 months of age identified TLR and DAP12 (TREM2 adaptor) signaling as the most significant pathway associated to gal3 in the 5xFAD mice. This gene expression data suggests an important role of gal3 in the AD neuroinflammatory response, perhaps by stimulating TLR4 and/or other TLRs [10, 33, 45] or glycoproteins, such as TREM2 [15, 74], which is implicated in AD-associated immune responses.

Consequently, we investigated whether gal3 interacts with TREM2, a key receptor suggested to drive the DAM phenotype. Our confocal microscopy study demonstrated a remarkable cellular co-localization of TREM2 and gal3 in microglial cells around Aβ plaques and a near 100% correspondence between both microglial markers, an indication that gal3 specifically labels DAM. STORM microscopy demonstrated that TREM2 and gal3 physically interact on microglial processes, most likely at the cell membranes of DAM microglia. We also incubated sections from 5xFAD/Gal3KO mice with recombinant gal3 to analyze the potential TREM2-binding of extracellular gal3. This assay demonstrated the ability of gal3 to selectively bind to TREM2^+^ microglia but not to homeostatic microglia. Direct interaction between gal3 CRD and TREM2 was demonstrated by fluorescent anisotropy with a Kd value of 450 ± 90 nM, in the same range as other preferred glycoproteins [12, 13]. The ability of gal3 to stimulate TREM2 was finally confirmed by a TREM2-DAP12 reporter cell line. Overall, we conclude that the switch from homeostatic microglia to DAM is accompanied by a significant upregulation of TREM2 and gal3, which seem to work in concert. We cannot exclude the possibility that gal3 interacts with different microglial receptors including TLR4 [10], IGFR [40], MerTK [53] and TREM2 (this study). In fact, because gal3 is relatively promiscuous in its interactions with glycoproteins, it may be behind the chronic and detrimental activation of microglia in AD. Regardless of this possibility, what our study demonstrates is that the pleiotropic activity of gal3 drives the amyloid-associated immune responses.

Additionally, our study has uncovered an unexpected role of gal3 as a powerful Aβ cross-seeding agent acting at nanomolar concentrations. We have previously demonstrated the ability of LPS-induced reactive microglia to release gal3[10, 53]. More recently, we have demonstrated a significant increase of gal3 levels in CSF from mice exposed to traumatic brain injury[73], an indication that gal3 is released by reactive microglia. In this study, we performed electron microscopy using immunogold gal3 staining to confirm that plaque-associated microglia is the main cellular phenotype expressing gal3 and that gal3 is present in the extracellular space and in Aβ fibrillar aggregates. This suggested the possibility that gal3 released from reactive microglia may seed Aβ aggregation. Indeed, pure gal3 accelerated the aggregation of pure Aβ at a ratio 1:10 000, a clear indication that it acts as an endogenous cross-seeding agent of Aβ fibrillation. Imaging and co-sediment experiments performed after incubations between Aβ and gal3 confirmed the amyloid aggregating effect of gal3.

To further demonstrate that gal3 acts as an Aβ cross-seeding agent *in vivo*, we injected Aβ monomers with or without gal3 into the hippocampi of WT mice; each side of the hippocampi was injected with either Aβ monomers alone or Aβ monomers and gal3. Aβ deposition was analyzed two months after injections. While no Aβ deposition was found in animals injected with Aβ monomers alone, co-injection of Aβ monomers and gal3 resulted in evident, insoluble Aβ aggregates, a clear demonstration that gal3 is acting as a strong Aβ cross-seeding agent. To induce Aβ aggregates by injecting Aβ species *in vivo*, an AD mouse model is typically a prerequisite[19]. Kim et al. injected Aβ peptides in normal mice, inducing AD-like cognitive deficits [36]. However, they did not demonstrate the formation of plaques in the injected area.

Recently, Heneka and colleagues have demonstrated that ASC specks released from reactive microglia physically interacts with Aβ, acting as an Aβ cross-seeding agent (Venegas et al., 2017). Eliminating microglia has been shown to prevent plaque formation in APP transgenic mice [Sosna et al 2018; PMID: 29490706], suggesting that factors released from microglia may seed amyloid plaques. Our study reinforces the view that reactive microglia play a critical role in Aβ aggregation and associated immune response by releasing Aβ cross-seeding agents (*i.e.* ASC specks and gal3). Aβ fibrillation is believed to precede the appearance of clinical symptoms and, hence, elucidating the earlier molecular mechanism involved in plaque formation appears critical for the establishment of promising therapeutic strategies aimed at stopping the development of AD at the first step. Gal3-inhibition emerges as a promising Aβ therapeutic target. The ability of gal3 to drive pro-inflammatory fAβ-associated immune responses, cross-seed Aβ and hinder Aβ clearance points (*i.e.* phagocytosis and IDE-1 levels) makes this protein a strategic upstream regulator of AD pathology. Indeed, the pathogenic role of gal3 was confirmed in adult 5xFAD mice lacking gal3 as those mice had a significantly lower Aβ load and ameliorated cognitive/spatial memory deficits as studied by the Morris water maze test.

Our results suggest that *LGALS3* gene variants affect the risk to develop AD, as indicated by the 5 SNPs in the *LGALS3* gene that we found to be related to increased AD frequency. However, according to the GWAS catalogue (https://www.ebi.ac.uk/gwas/), none of the variants reported in Suppl. Table 3 or those in linkage disequilibrium with them (Suppl. Fig. 3B), have been previously associated with AD. This could be due to the fact that the GWAS approach requires large samples to detect modest effects, partly due to the multiple testing corrections applied. However, the impact of these *LGALS3* SNPs appear to be similar to other SNPs included in the GWAS, supporting a role of this gene in the development of AD. Our meta-analysis only comprised those SNPs belonging to a linkage disequilibrium block located at the 5’ end of *LGALS3* gene, and, therefore, we do not know if other regions could have genetic variants showing stronger effects. Further replication studies covering the entire genetic region of this locus will be necessary to confirm our results.

## CONCLUSIONS

In conclusion, we provide evidence that gal3 is a central upstream regulator of microglial immune response in AD. It drives pro-inflammatory activation of microglia in response to fAβ along with impairment of fAβ degradation and clearance. Our study has additionally uncovered two exciting features of gal3 disease biology that appears important in AD pathogenesis. First, gal3 is an endogenous TREM2 ligand, a key receptor driving microglial activation in AD. Second, nanomolar concentrations of gal3 cross-seeds Aβ aggregation *in vitro* and *in vivo.* As a result, gal3 inhibition may be a potential pharmacological approach to counteract AD at multiple stages of the disease.

## Supporting information

## Author contribution

A.B-S., T.D., R.R., R.S-V., A.P., I.F-J., Y.Y., M.W., A.V., D.A., C.D., J.S., S.J., V.N-G., J.G., L.M.R., and S.L. performed experiments and analyzed data. M.W. and E.E. provided human samples. A.B-S., G.C.B., T.D., J.L.V and J.V. were involved in the study design and wrote the paper. All authors discussed results and commented on the manuscript.

## Acknowledgement

This work was supported by grants from the Swedish Research Council, Bagadilico (Linné consortium sponsored by the Swedish Research Council) and the Strong Research Environment MultiPark (Multidisciplinary Research in Parkinson’s and Alzheimer’s Disease at Lund University), the Swedish Alzheimer’s Foundation, Swedish Brain Foundation, A.E. Berger Foundation, Gyllenstiernska Krapperup Foundation, the Royal Physiographic Society, Crafoord Foundation, Olle Engkvist Byggmästare Foundation, Wiberg Foundation, G&J Kock Foundation, Stohnes Foundation, Swedish Dementia Association and the Medical Faculty at Lund University. This work was supported by grant SAF2015-64171R (Spanish MINECO/FEDER, UE), Fondo de Investigación Sanitaria (FIS), projects FIS PI12/O1431 and FIS PI15/00796, from Instituto de Salud Carlos III (ISCiii) of Spain, co-financed by FEDER funds from European Union, through grants PI12/01439, PI15/00957 (to JV) and PI12/01431, PI15/00796 (to AG), and by Junta de Andalucia, Proyecto de Excelencia (CTS-2035) (to JV and AG). AV and GCB received funding from the Innovative Medicines Initiative 2 Joint Undertaking under grant agreement No 115976 (PHAGO). CIBERNED “Centro de Investigacion Biomedica en Red sobre Enfermedades Neurodegenerativas (CIBERNED), Madrid (Spain)”. HL, AF was supported by the Swedish Research Council, the Swedish Brain Foundation, the Alzheimer Foundation and the Åhlén Foundation. UJN was supported by grants from the Knut and Alice Wallenberg Foundation (KAW 2013.0022) and the Swedish Research Council (grant no. 621-2012-2978). We thank Barbro-Kahl Knutson for help with producing and analyzing galectins, and Fredrik Zetterberg and Hans Schambye at Galecto Biotech AB, Sweden, for providing galectin inhibitors.

---

**Suppl. Fig. 1.**
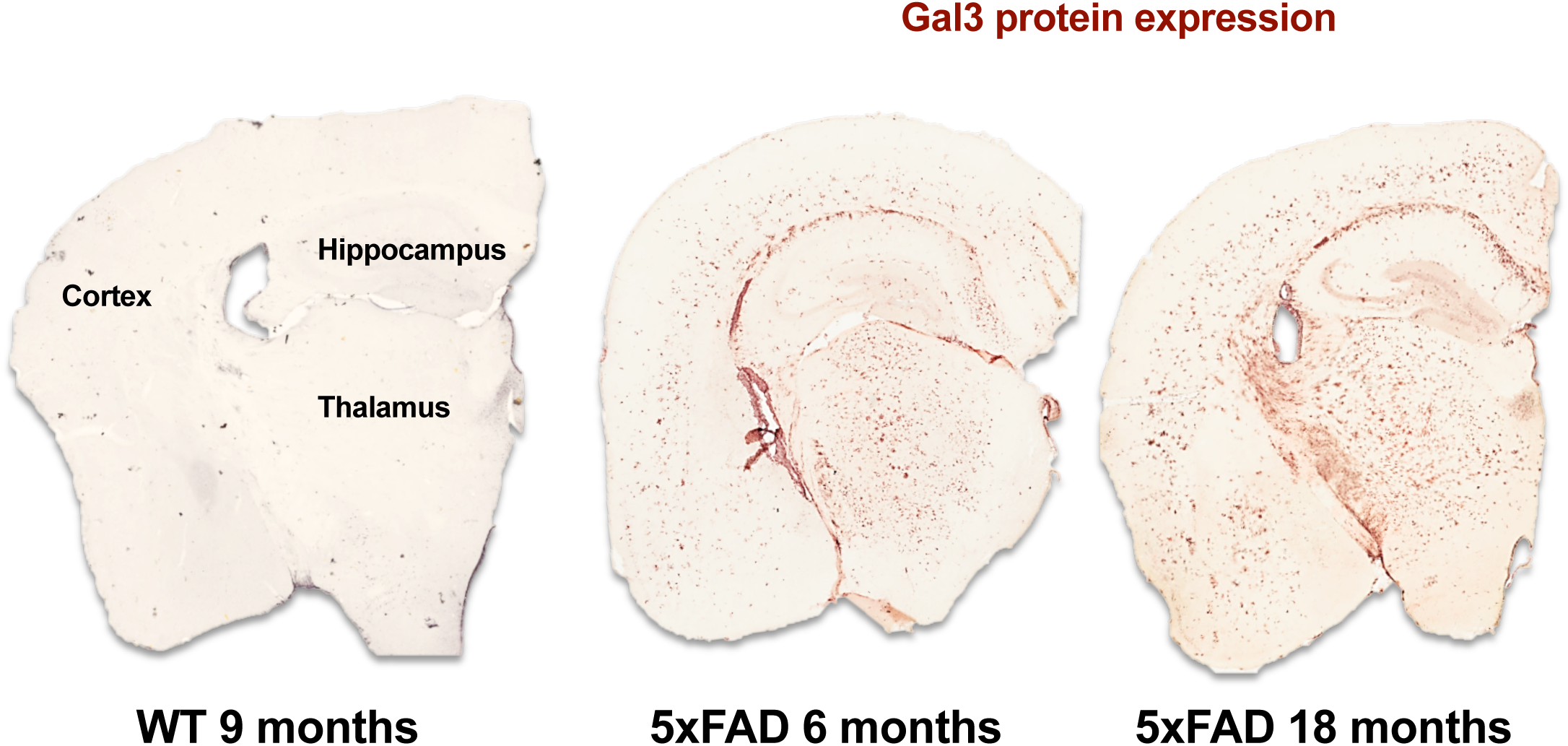

**Suppl. Fig. 2.**
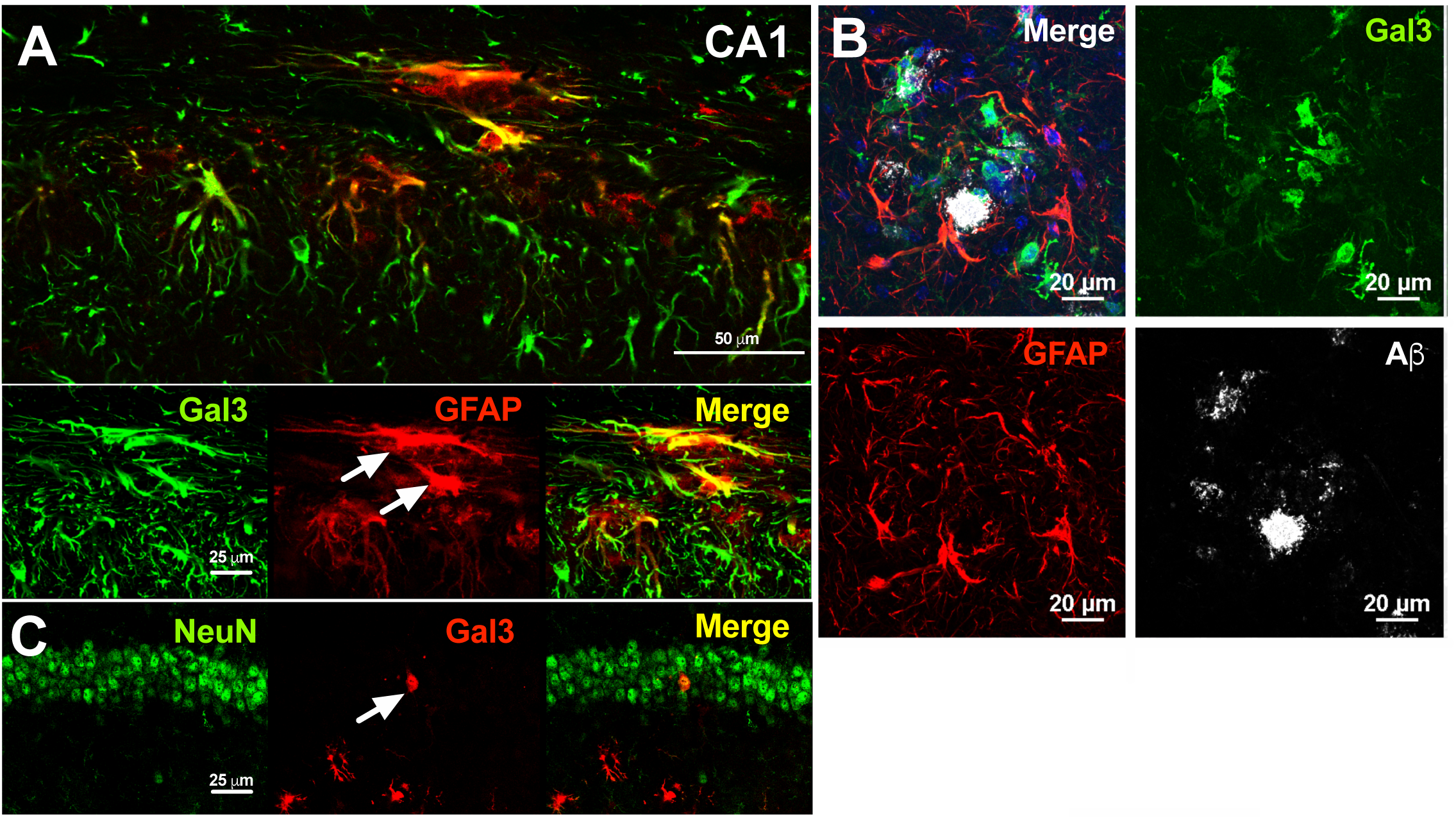

**Suppl. Fig. 3.**
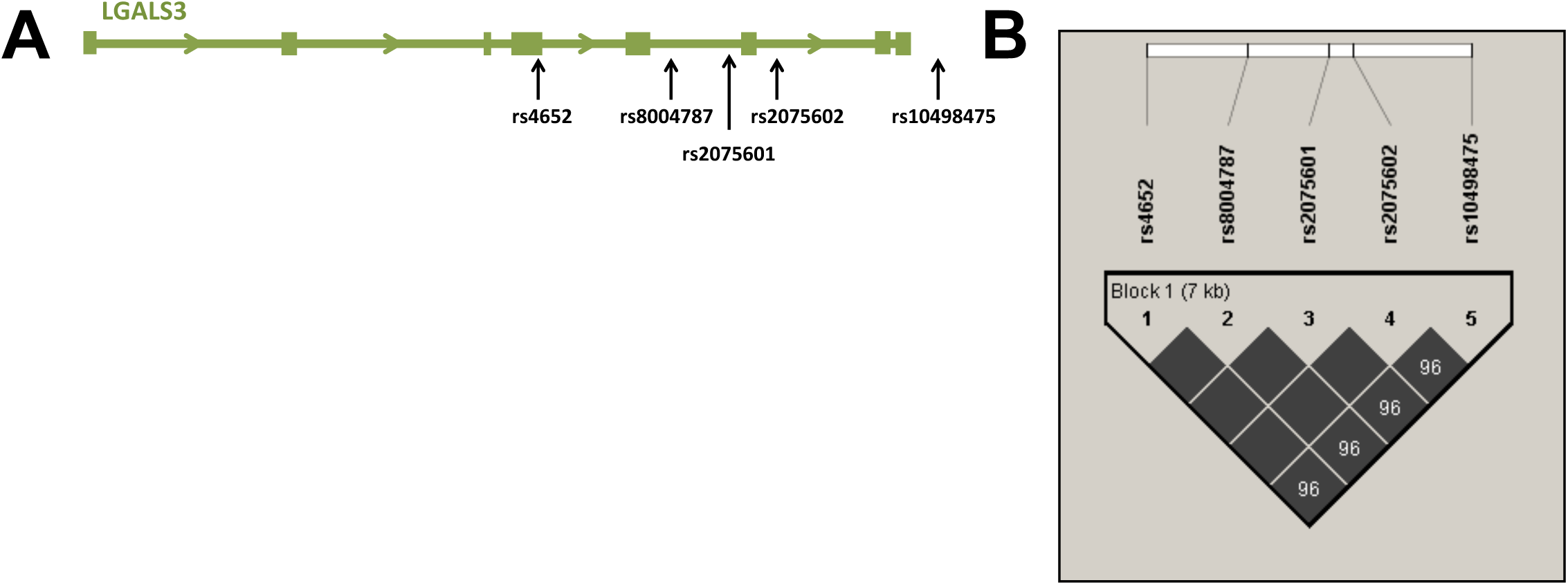

**Suppl. Fig. 4.**
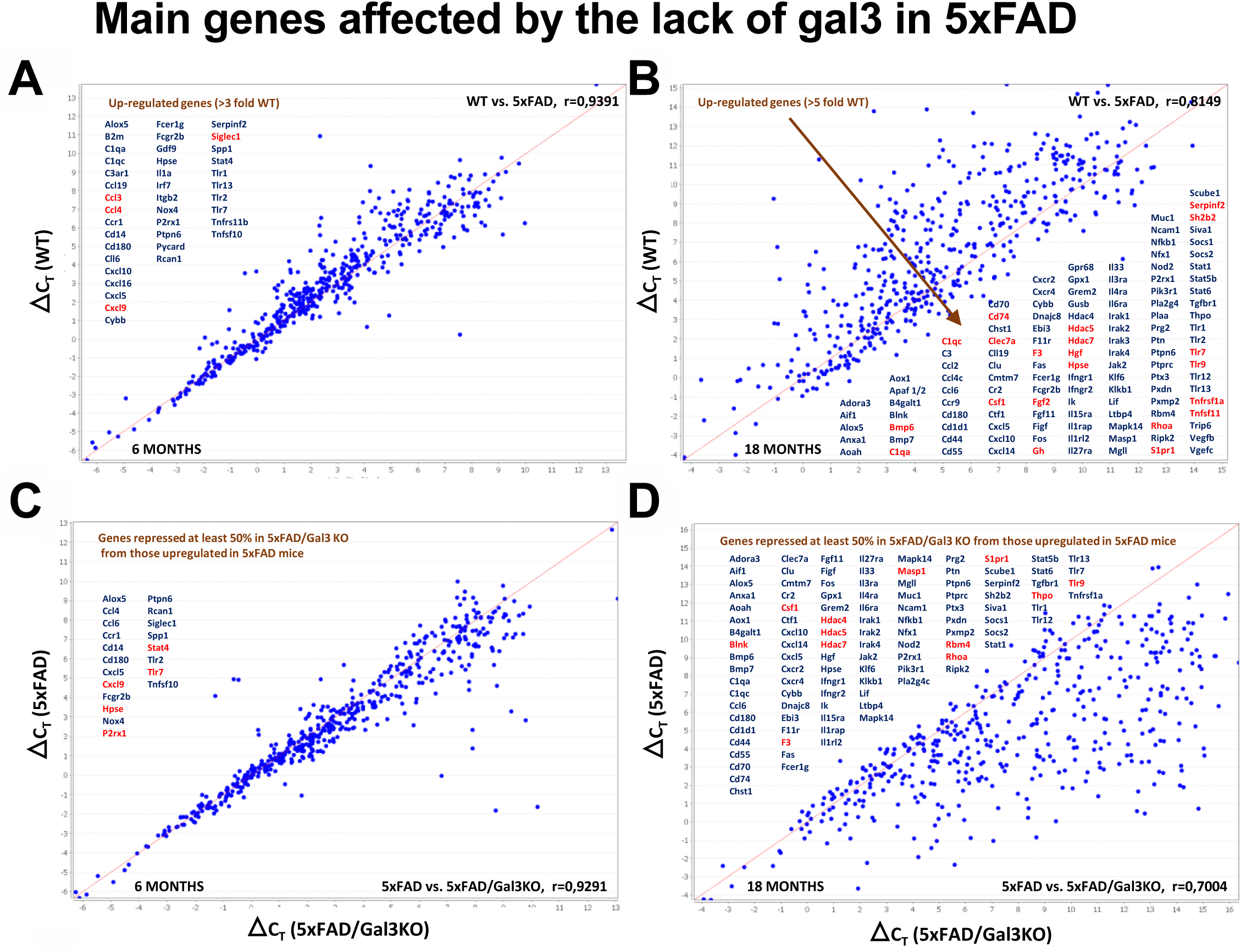

**Suppl. Fig. 5.**
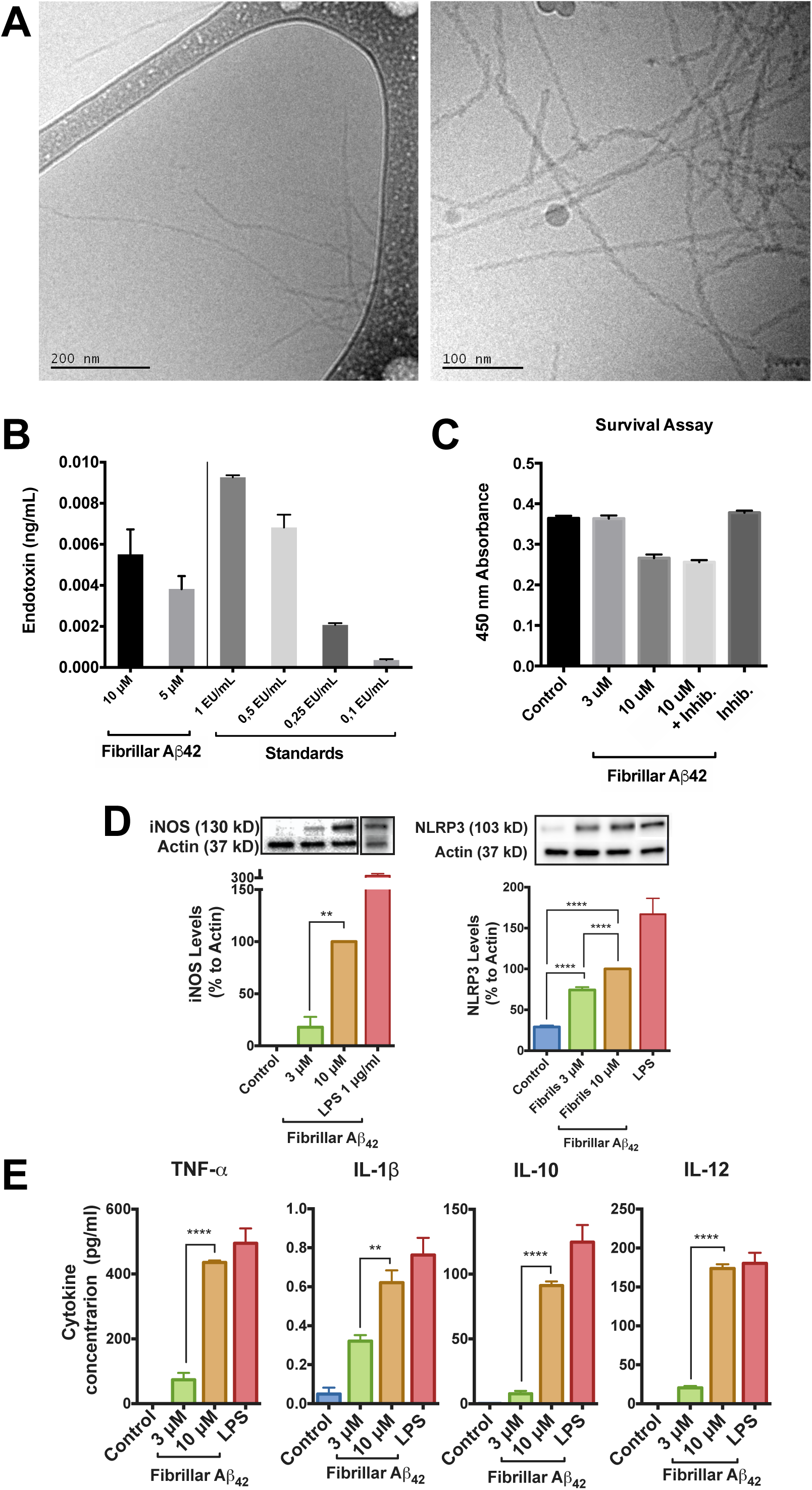

**Suppl. Fig. 6.**
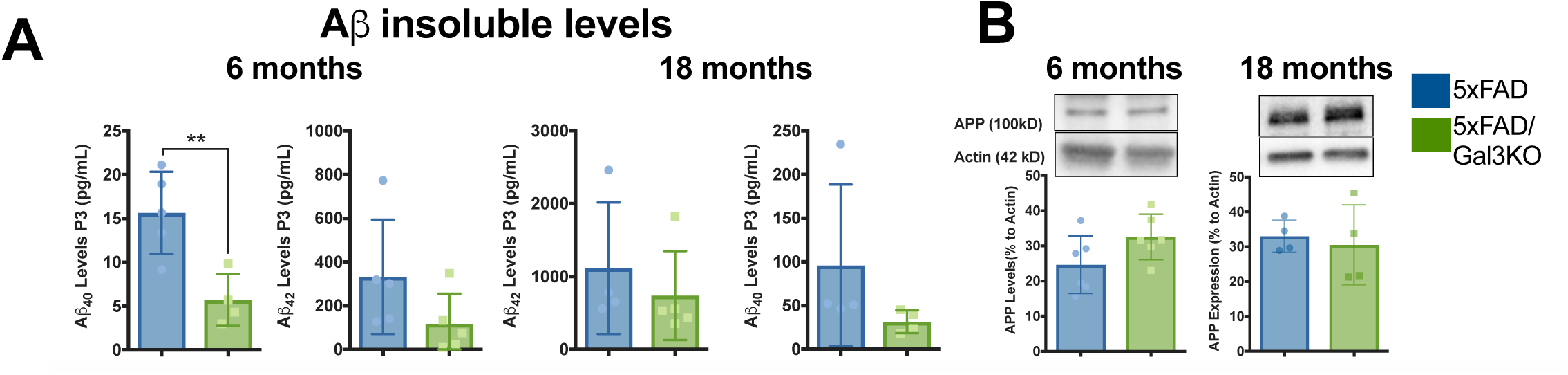

**Suppl. Fig. 7.**
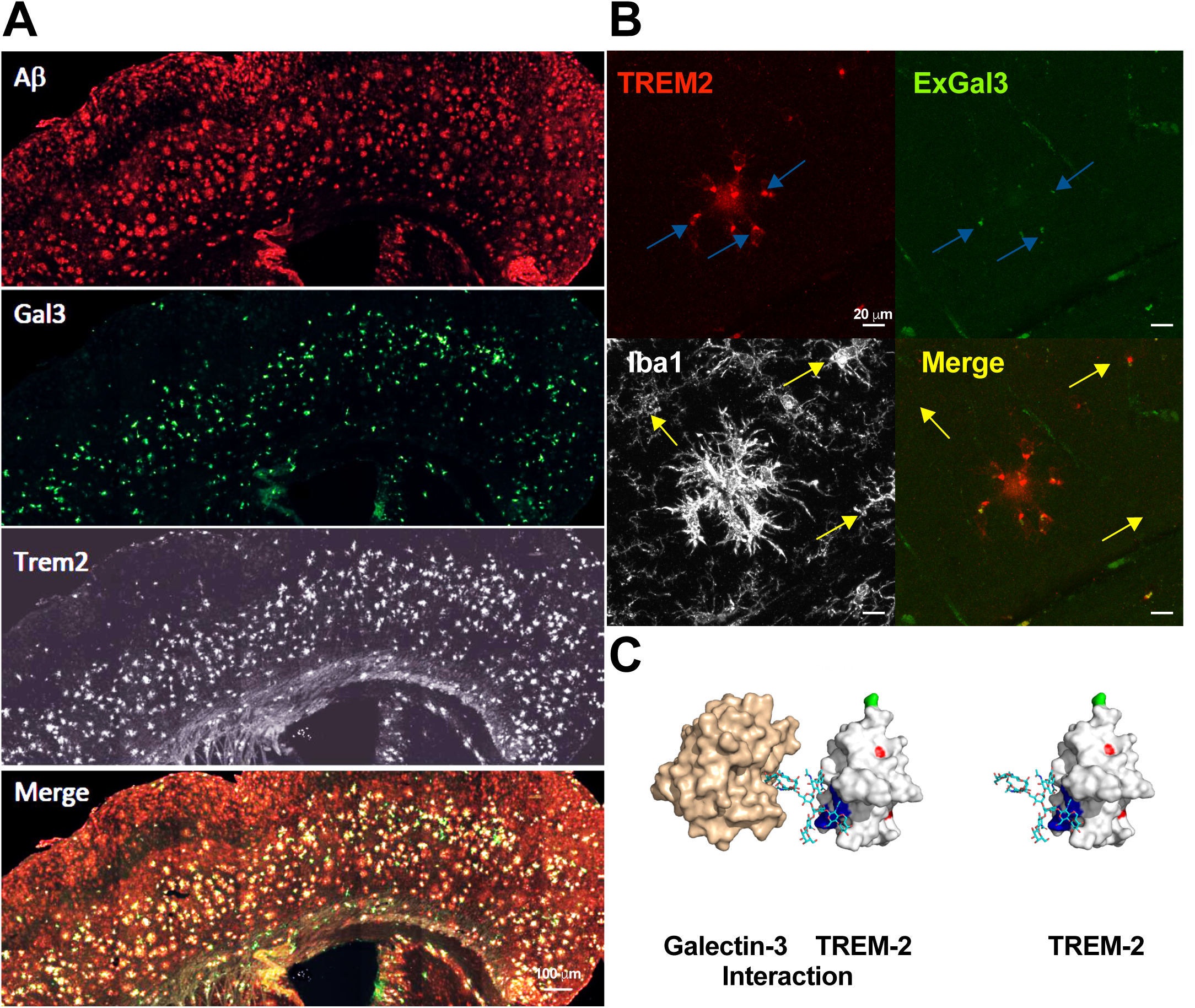

