## Supplementary material for "Galectin-3, A Novel Endogenous Trem2 Ligand, Regulates Inflammatory Response and Aβ Fibrilation in Alzheimer’s Disease"

| CHR | BP | SNP | A1 | A2 | N | P | P(R) | OR | OR(R) | Q | I | ADNI | INIA | GENADA | MUR |
|---|---|---|---|---|---|---|---|---|---|---|---|---|---|---|---|
| 14 | 54674789 | rs4652 | C | A | 3 | 0.022 | 0.022 | 1.11 | 1.11 | 0.4455 | 0.00 | 1.33 | 1.13 | 1.07 | NI |
| 14 | 54677119 | rs8004787 | T | C | 3 | 0.027 | 0.023 | 1.10 | 1.11 | 0.4418 | 0.00 | 1.33 | 1.10 | 1.07 | NI |
| 14 | 54678987 | rs2075601 | T | C | 3 | 0.022 | 0.022 | 1.11 | 1.11 | 0.4455 | 0.00 | 1.33 | 1.11 | 1.07 | NI |
| 14 | 54679527 | rs2075602 | G | A | 3 | 0.027 | 0.023 | 1.10 | 1.11 | 0.4418 | 0.00 | 1.33 | 1.10 | 1.07 | NI |
| 14 | 54682233 | rs10498475 | T | C | 3 | 0.018 | 0.018 | 1.27 | 1.23 | 0.4896 | 0.00 | 0.96 | 1.25 | NI | 1.51 |

CHR, chromosome; BP, base pair position; SNP, single nucleotide polymorphism; A1, reference allele; A2, alternative allele; N, number of GWAS included in the meta-analysis; p, fixed-effects p-value; p (R), random-effects p-value; OR, fixed-effects odds ratio; OR (R), random-effects odds ratio; Q, p-value for heterogeneity of OR; I, effect size for heterogeneity of OR. The last four columns show the OR for each GWAS.
