## Supplementary material for "Galectin-3, A Novel Endogenous Trem2 Ligand, Regulates Inflammatory Response and Aβ Fibrilation in Alzheimer’s Disease"

Table 1A. **AD subjects and healthy controls used to evaluate the role of galectin-3 in Alzheimer's disease.**

| *Case* | *Sex* | *Age (years)* | *Post Mortem Time* | *Cause of Death* | *CNS Diagnose* |
|---|---|---|---|---|---|
| Healthy Control 1 | Male | 80 | 5h 25min | Euthanasia (colon carcinoma) | - |
| Healthy Control 2 | Female | 68 | 6h 50min | Euthanasia (metastatic ovarian cancer and ileus) | - |
| Healthy Control 3 | Male | 65 | 9h 10min | Euthanasia (heart failure) | - |
| AD Patient 1 | Female | 64 | 5h 15 min | Pneumonia and stomach bleeding | Alzheimer's Disease |
| AD Patient 2 | Male | 64 | 8h 15 min | Euthanasia (AD with Lewy's bodies) | Alzheimer's Disease with Lewis Bodies |
| AD Patient 3 | Female | 72 | 4h 25 min | Advanced Alzheimer's with delirium | Advanced Alzheimer's with delirium |
| AD Patient 4 | Male | 64 | 6h 40 min | Dehydration after CVA and AD | Alzheimer's Disease |

Patients and healthy controls used to evaluate the presence of Gal3 in the vicinity of the senile plaques by immunohistochemistry.

Table 1B. **AD subjects and healthy controls used to evaluate the role of galectin-3 in Alzheimer's disease.**

| ***Case*** | ***Sex*** | ***Age (years)*** | ***Post Mortem Time*** | ***Cause of Death*** | ***CNS Diagnose*** |
|---|---|---|---|---|---|
| Healthy Control 1 | Male | 80 | 72h | AMI | - |
| Healthy Control 2 | Male | 68 | 24h | Cardiac Insuff | - |
| Healthy Control 3 | Female | 65 | 24h | AMI | - |
| Healthy Control 4 | Female | 86 | 4 days | Aorta-diss. O rupture | - |
| Healthy Control 5 | Male | 75 | 72h | AMI | - |
| AD patient 1 | Male | 77 | 48h | Pneumonia | Alzheimer's Disease-FTD |
| AD Patient 2 | Male | 82 | 7 days | AMI+Septicemia | Alzheimer's Disease |
| AD Patient 3 | Male | 63 | 96h | Pulm Embolus | Alzheimer's Disease |
| AD Patient 4 | Male | 76 | 48h | Pneumonia + AMI | Alzheimer's Disease |
| AD Patient 5 | Male | 80 | 48h | Pneumonia + Car.Insuff | Alzheimer's Disease |
| AD Patient 6 | Female | 70 | 4 days | Pneumonia + AMI | Alzheimer's Disease |

*AMI = *Acute Myocardial Infarction*. Patients and healthy controls were used to evaluate the inflammatory component (IL6), the levels of Gal3 and amyloid beta by western blot and ELISA.
